# Biochemically-functionalized probes for cell type-specific targeting and recording in the brain

**DOI:** 10.1101/2023.10.02.560579

**Authors:** Anqi Zhang, Theodore J. Zwang, Charles M. Lieber

**Affiliations:** Department of Chemical Engineering and Department of Bioengineering, Stanford University; Stanford, California, 94305, USA; Department of Chemistry and Chemical Biology, Harvard University; Cambridge, Massachusetts 02138, USA; MassGeneral Institute for Neurodegenerative Disease, Massachusetts General Hospital; Boston, Massachusetts, 02114, USA; Department of Neurology, Harvard Medical School; Boston, Massachusetts, 02114, USA

## Abstract

Selective targeting and modulation of distinct cell types and neuron subtypes is central to understanding complex neural circuitry, and could enable electronic treatments that target specific circuits while minimizing off-target effects. However, current brain-implantable electronics have not yet achieved cell-type specificity. We address this challenge by functionalizing flexible mesh electronic probes, which elicit minimal immune response, with antibodies or peptides to target specific cell markers. Histology studies reveal selective association of targeted neurons, astrocytes and microglia with functionalized probe surfaces without accumulating off-target cells. In vivo chronic electrophysiology further yields recordings consistent with selective targeting of these cell types. Last, probes functionalized to target dopamine 2 receptor expressing neurons show the potential for neuron subtype specific targeting and electrophysiology.

## Introduction

A central step in deciphering complex brain circuits is to identify the roles of specific cell types and neural subtypes (*1-4*). The ability to target specific cell types for recording and modulation is critical to achieving this goal. With the advent of optogenetics (*5*), for example, specific cell types can be engineered such that they express light-sensitive channelrhodopsins (*6*), thereby enabling targeted optogenetic stimulation. However, cell-type specific recording and modulation has not been possible with implantable electrical probes. Traditional probes made of silicon, for example, have substantially higher stiffness than soft brain tissue and larger feature size than individual neurons, which can lead to chronic immune response including build-up of glial scar tissue and depletion of neurons immediately adjacent to electrodes (*7, 8*). Considerable effort has been placed on increasing substrate flexibility and reducing feature size (*7-11*) to address the chronic immune response. For example, mesh electronic probes (*12-14*), which were designed with tissue-like flexibility and a macroporous structure, do not elicit inflammatory immune response and show little effect on the natural distribution of neurons and glial cells postimplantation. The minimal immune response makes them an attractive substrate for functionalization with antibodies or peptides targeting specific cell types or neural subtypes, which might allow for selective *in vivo* electrophysiology without genetic modification.

Previously, surface modification approaches have been utilized to functionalize the surfaces of chip-based electrodes and nanoprobes for cellular measurements (*15-19*). For example, attachment of RGD-based peptides that trigger adhesion and engulfment mechanisms have been shown to yield intracellular-like recording with mushroom-like gold electrodes (*16*), while phospholipid layers on the surfaces of nanoscale transistor probe arrays facilitated internalization and intracellular recording in primary neurons and cardiac cells (*17-19*). In addition, substantial work has been carried out functionalizing the surfaces of implanted neural probes with biologically-derived or biomimetic materials (*20, 21*), including neuronal cell adhesion molecules, extracellular matrix molecules, and conductive polymers. These biocompatible surface coatings effectively reduced the immune response and promoted neuronal survival adjacent to the neural probes, but have not shown the capability to target and monitor specific cell types.

## Results

### Implantable mesh probes with cell type specificity

Mesh electronic probes were first modified with either a control antibody or antibodies and peptides that can recognize specific surface receptors of different cell types, including neurons, astrocytes and microglia (Fig. 1A). To target neurons, we selected the peptide sequence Ile-Lys-Val-Ala-Val (IKVAV), a fragment of laminin α-1 that has been demonstrated to promote neuronal cell attachment on substrates while minimizing glial cell adhesion (*22-24*). To target astrocytes, we chose an antibody targeting the extracellular domain of excitatory amino acid transporter 2 (EAAT2), a membrane-targeted glutamate transporter highly expressed in mature astrocytes (*25*). To target microglia, we chose an antibody targeting the microglia surface marker CD11b (*26*). To exclude the possible effects of immune response (i.e., accumulation of astrocytes and microglia) induced by foreign antibodies anti-EAAT2 and anti-CD11b, we included a rabbit anti-human immunoglobulin G (IgG) antibody as a negative control (control Ab), because it was produced in the same host species and does not specifically bind to cells in the mouse brain.

**Fig. 1.**
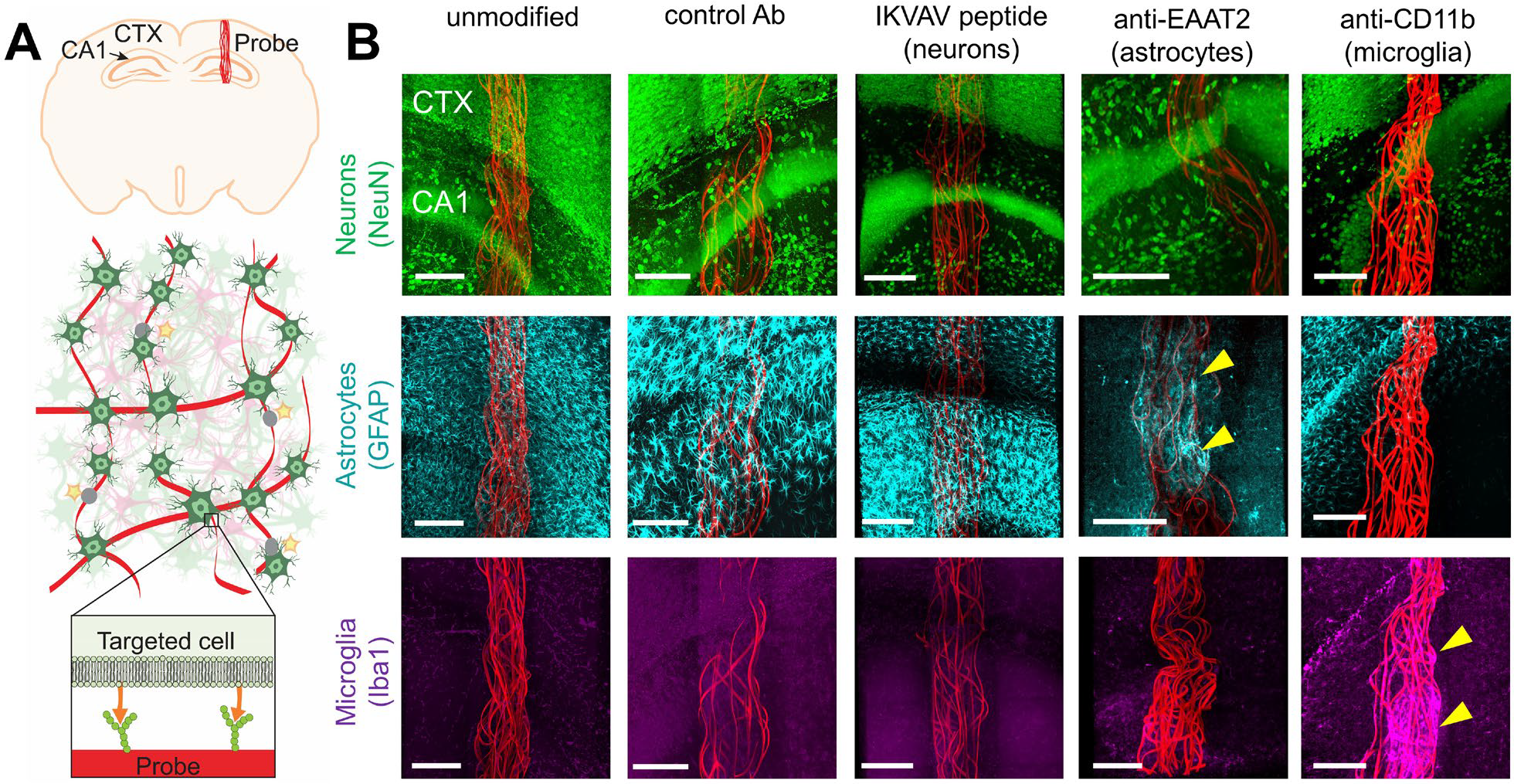
Implanted control and functionalized mesh probes. (**A**) Top, schematic of a coronal brain section showing a mesh probe (red) implanted through the cortex (CTX) into the mouse hippocampus with the cornu ammonis (CA1) region indicated. Bottom, schematics of small volumes of brain tissue/probe interface highlighting probes functionalized with antibodies and peptides capable of targeting specific cell types. Mesh probe ribbons are in red. Cells in green are targeted cells; cells in pink are non-targeted cells. (**B**) Representative fluorescence images of coronal mouse brain section of control and functionalized probes including the hippocampal CA1 region 14 days post-implantation. The five columns correspond to an unmodified probe as well as four distinct probes functionalized with either control antibody (Ab, IgG), IKVAV peptide for targeting neurons, anti-EAAT2 Ab for targeting astrocytes or anti-CD11b Ab for targeting microglia. The three rows highlight the tissue staining of the section (across the five distinct probe modifications) with NeuN, GFAP and Iba1 antibodies for neurons, astrocytes, and microglia, respectively. The yellow triangles in the anti-EAAT2-GFAP and anti-CD11b-Iba1 panels highlight the specific targeting of astrocytes and microglia, respectively. Scale bars, 200 µm.

The probes (fig. S1) consist of double-sided platinum electrodes (*27*) embedded in the polymer-based mesh-like region and connected through a stem to the input/output (I/O) structure that provides an electrical interface for external recording as described previously (*28*). Surface functionalization of the mesh region is based on standard covalent coupling chemistry (*29*) that uses the carboxyl (-COOH) groups on the surface of the SU-8 polymer resulting from the oxygen plasma treatment at the end of probe fabrication (*30*). Previous studies have shown that this covalent coupling chemistry should yield at least 3-month stability under conditions relevant to our experiments (*31*). The modification process was carried out by first loading probes into glass capillary tubes, and then dipping the end closer to the mesh region into the respective modification solutions (see materials and methods). The primary amine (-NH_2_) groups on the peptides and primary antibodies are covalently-coupled to the -COOH groups on the probe surface using 1-ethyl-3-(3-(dimethylamino)propyl) carbodiimide (EDC)/N-hydroxysuccinimide (NHS) (*29*). The effectiveness of the coupling was characterized by confocal fluorescence microscopy where either fluorophore-labeled IKVAV peptides were directly coupled or secondary antibodies incubated with the conjugated primary antibodies (fig. S2). The images show uniform fluorescence on the SU-8 polymer ribbons consistent with specific modification of the probe surface without aggregates of either the peptide or antibodies layers.

The unmodified and functionalized probes were implanted into the mouse hippocampus using a stereotaxic injection method (*32*) (Fig. 1A). 14 days after implantation, the brains were extracted, sectioned, cleared, and stained using a modified CLARITY (*33*) protocol (see materials and methods) to allow fluorescence imaging of the cellular environment around the intact probe. We labeled the brain tissue with antibodies for neurons (anti-NeuN), astrocytes (anti-GFAP) and microglia (anti-Iba1) and used confocal microscopy to image the hippocampal CA1 region in 1-mm thick tissue sections containing the intact probes. We observed striking differences in the cellular environments near probes with different modifications (Fig. 1B). For unmodified, control Ab-modified, and IKVAV peptide-modified mesh probes, neuron somas and axons fully penetrated the probe ribbons, with neurons, astrocytes and microglia exhibited similar distribution inside and outside the mesh boundaries. The anti-EAAT2- and anti-CD11b-modified mesh probes, however, showed significantly enhanced accumulation of astrocytes and microglia, respectively. Qualitative comparison of how the different probe surfaces affected cell distributions reveals several important points. First, the similar cell distribution near unmodified, control Ab, and peptide-modified mesh probes indicates that antibodies produced in different species (e.g., rabbit) did not cause a substantial immune response within the observed 14-day time frame, and that the modification strategy did not have an adverse effect on the intrinsic biocompatibility of the mesh probes. Second, the anti-EAAT2 and anti-CD11b probes selectively attracted astrocytes and microglia, respectively, without affecting off-target cell types, which confirmed our initial hypothesis that the surface functionalization with specific antibodies could selectively recruit different cell types. Third, the accumulated cells were only observed near the surface of the mesh ribbons, indicating that the modifications only affected cells close to the functionalized probes.

We then asked whether quantitative analyses of modified probes implanted into mouse brains could provide more insight into these qualitative observations as well as the potential for recruitment of neurons by examination of time-dependent histology data (Fig. 2, figs. S3 - S9). Histology imaging was replicated on samples (at least N = 3 mice at each time point) containing mesh probes with different modifications on days 3, 7, and 14 post-implantation (Fig. 2A-C, fig. S3, fig. S4). We identified individual neurons, astrocytes, and microglia in the CA1 region using reported methods (*34, 35*), and then plotted the different cell density distributions up to 300 μm away from the probe surface (see materials and methods, Fig. 2D-F, fig. S3, fig. S4). The cell density was compared quantitatively by binning all cells at a given distance from the probe surface. Each value from a given sample was normalized to the average value from all bins in that sample.

**Fig. 2.**
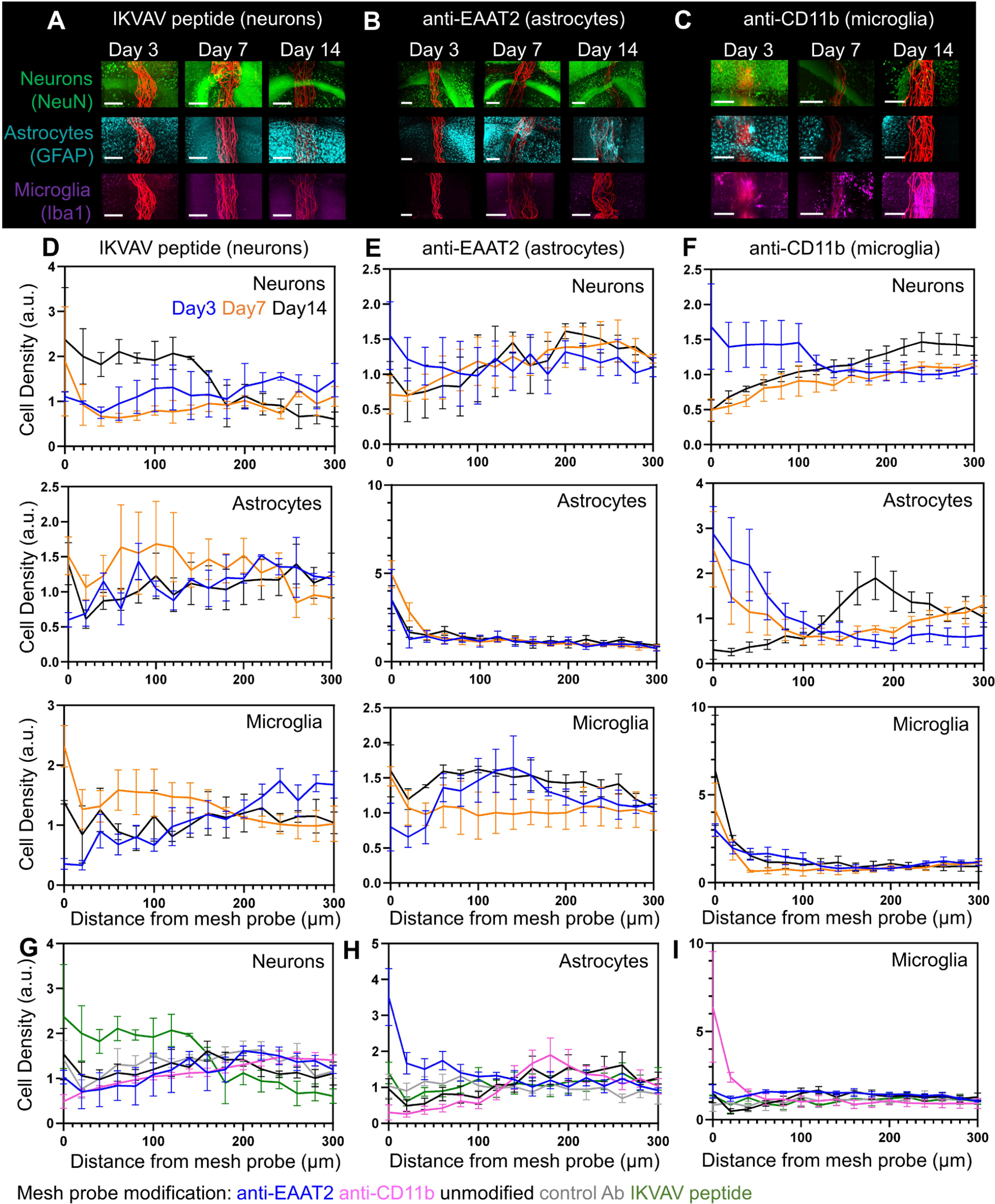
Time-dependent histology studies of mesh probe–brain interfaces. (**A**)-(**C**) Time-dependent confocal microscopy images of cleared tissues slices at 3, 7, and 14 days post-implantation labeled with NeuN (neurons, top), GFAP (astrocytes, middle), and Iba1 (microglia, bottom) for mesh probes functionalized with (A) IKVAV, (B) anti-EAAT2, and (C) anti-CD11b. Scale bars, 200 µm. (**D**)-(**F**) Normalized cell density distribution for neurons (top), astrocytes (middle), and microglia (bottom) in the CA1 region as a function of distance from surface of mesh probes modified with (D) IKVAV peptide, (E) anti-EAAT2, and (F) anti-CD11b on days 3 (blue), 7 (orange), and 14 (black) (at least N = 3 mice at each time point). (**G**)-(**I**) Normalized cell density distribution for (G) neurons, (H) astrocytes, and (I) microglia as a function of distance from the probes modified with anti-EAAT2 (blue), anti-CD11b (pink), unmodified (black), control antibody (grey), and IKVAV peptide (green) 14 days post-implantation. Values are means ± standard error of the mean (s.e.m.). See figs. S5-S9 for cell density distribution plots for individual samples.

Consistent with our previous observations, the unmodified (*14*) (fig. S3) and control Ab-modified mesh probes (fig. S4) only showed a small displacement of neurons near the probe surface that recovered over time, and no significant influence on the distribution of astrocytes or microglia. Notably, the other probes showed different time courses of cellular accumulation depending on their surface modifications. For IKVAV peptide-modified mesh probes (Fig. 2D), the neuron density increased 14 days post-implantation while the density of astrocytes and microglia always remained consistent with the baseline as with the control Ab. For probes modified with anti-EAAT2 (Fig. 2E), an increased density of astrocytes was observed at all time points near the mesh surface, while the microglia density remained close to the baseline level and the neuron density slightly decreased approaching the probe surface. For probes modified with anti-CD11b (Fig. 2F), there was an accumulation of microglia near the probes at all time points, the astrocyte density temporarily increased on day 3 and dropped below the baseline on day 14, and the neuron density decreased when approaching the probe surface. A summary of the distributions of neurons, astrocytes, and microglia adjacent to mesh probes with different modifications 14 days post-implantation (Fig. 2G-I) further highlights the specific targeting of neurons, astrocytes and microglia by the IKVAV peptide, anti-EAAT2 and anti-CD11b-modified mesh probes, respectively. These observations demonstrate two important points. First, the specific accumulation of microglia and astrocytes occurred at a faster rate than that of the neurons, consistent with the higher mobility and enhanced migration of microglia and astrocytes than neurons after acute brain injury (*36*). Second, during recovery, astrocytes and microglia would selectively accumulate near the probe surface depending on the modification, suggesting the process was driven by modification rather than generalized inflammation where both cell types are expected to accumulate (*36*).

### Cell type-specific electrophysiological recording

Next, we asked how the specific targeting of different cell types affects electrophysiology recording. Since the neuron spike amplitude recorded by extracellular electrodes decreases exponentially with distance from the electrode (*37*), we hypothesized that IKVAV peptide-modified probes should show an increase in the number of recorded neurons and increase in mean spike amplitudes, while anti-EAAT2- and anti-CD11b-modified mesh probes would show a decrease in both quantities, compared to unmodified and control Ab-modified mesh probes.

Probes with different surface functionalization (N = 3 mice for each condition) were implanted through the cortex and into the hippocampus of mice for *in vivo* chronic recording measurements. Multiplexed recordings from head-fixed, awake mice (Fig. 3A-C, fig. S10) exhibited stable single-unit spikes from the time of initial implantation to the endpoint of the measurements (day 1 to day 14, fig. S11, fig. S13, fig. S15, fig. S17, fig. S19). Single neuron activity was tracked by clustering the sorted spikes with principal component analysis from all channels as a function of time (*12*) (figs. S12, fig. S14, fig. S16, fig. S18, fig. S20). These recordings highlight several key points. First, the IKVAV peptide-modified and unmodified mesh probes showed more single-unit spikes than the anti-CD11b and anti-EAAT2 mesh probes. Second, the control Ab-modified probes showed a similar level of spontaneous firing to that recorded by the unmodified probes. Third, the number of distinct neurons detected, represented by different color waveforms, increased mainly within the first 7 days, consistent with our previous results with unmodified mesh probes (*14*).

**Fig. 3.**
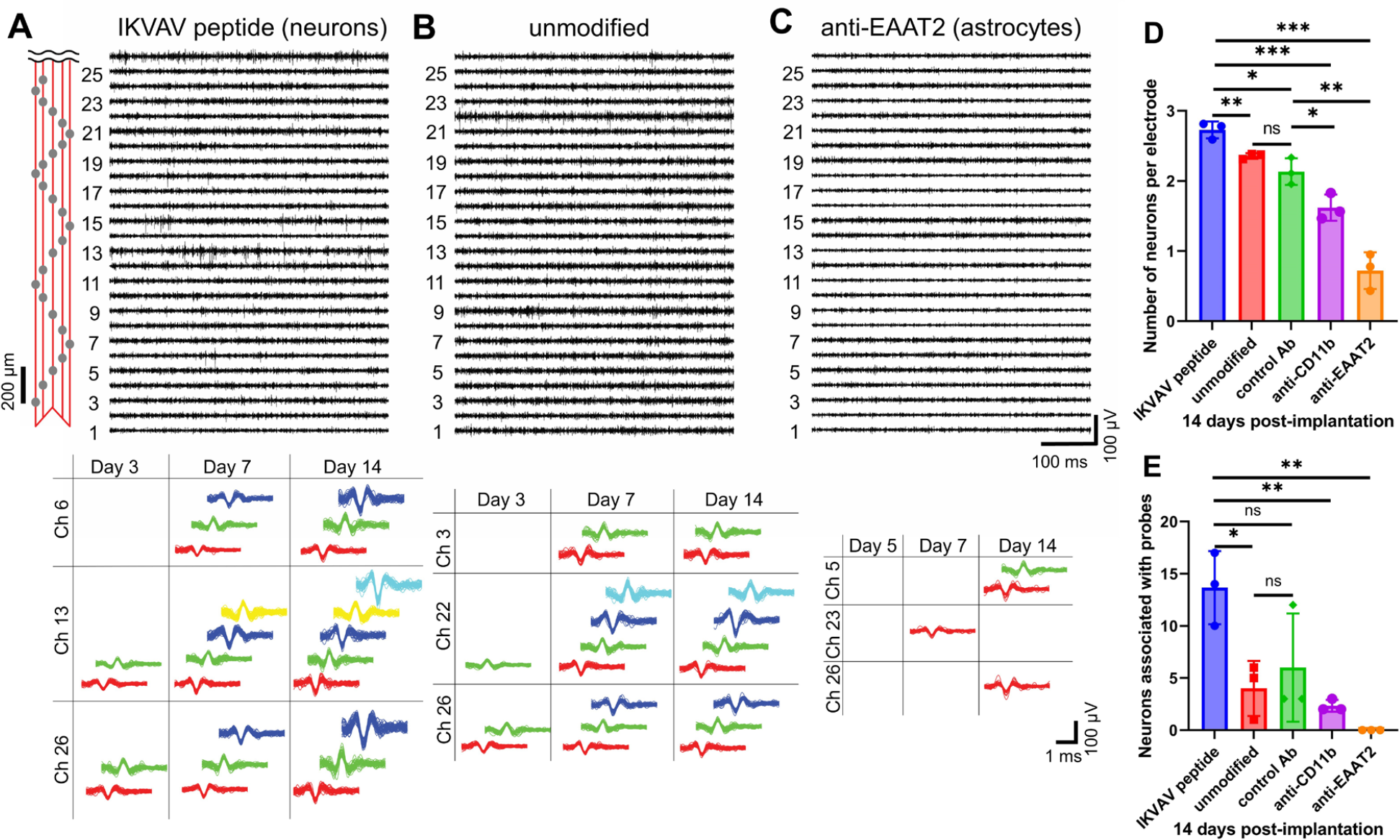
Cell type-specific electrophysiological recording and analysis. (**A**)-(**C**) Top, representative 26-channel single-unit spike traces 14 days post-implantation from mesh probes modified with IKVAV peptide for targeting neurons (A), unmodified (B), and modified with anti-EAAT2 for targeting astrocytes (C). Recordings were made from head-fixed, awake mice. The x and y axes represent recording time and voltage. The relative positions of the recording electrodes are marked by grey dots in the schematic (leftmost panel). Channels with smaller numbers are implanted deeper. Bottom, time evolution principal component analysis clustered single unit spikes from three representative channels. For each channel, each distinct color in the sorted spikes represents a unique identifiable neuron. (**D**) Average number of distinct neurons recorded per electrode 14 days post-implantation from mesh probes with different modifications (N = 3 mice). (**E**) Number of detected neurons closely associated with probes for the different modifications 14 days post-implantation (N = 3 mice). Close association of recorded neurons is defined as amplitude greater than mean amplitude of all mesh probes + 1 s.d.. Values are means ± s.d., ns nonsignificant, * P ≤ 0.05, ** P ≤ 0.01, *** P ≤ 0.001, two-tailed, unpaired, t-test.

We carried out statistical analyses of the sorted spikes from the probes with different modifications (Fig. 3D-E, fig. S21) to better understand these results. The average numbers of distinct neurons recorded per electrode from probes with different modifications (Fig. 3D) showed a significant increase for the IKVAV peptide-modified probes and a significant decrease for the anti-CD11b- and the anti-EAAT2-modified probes. In addition, the control Ab-modified probes appear to have recorded similar numbers of neurons as the unmodified probes. To specifically examine the number of neurons closely associated with the probe surface, we compared the amplitudes of all recorded neurons, and set the amplitude threshold for close association as the mean + s.d. (49.5 + 14.3 mV), since the spike amplitude decreases in an exponential manner as the distance from the electrode increases (*37*) (Fig. 3E). 14 days postimplantation, the IKVAV peptide-modified mesh probes recorded significantly higher numbers of closely associated neurons, while both anti-CD11b and anti-EAAT2 mesh probes showed significantly lower numbers of such neurons. These analyses are consistent with our quantitative histology data that showed IKVAV peptide functionalized probes enhanced neuron density close to the surface, while the accumulation of microglia and astrocytes excluded neurons adjacent to the anti-CD11b and anti-EAAT2 modified probes. The similarity of electrophysiology results between unmodified and control Ab mesh probes further supports our assessment that the antibody modification itself does not provoke an immune response, and that antibodies are changing the cell distribution by associating with their respective membrane surface antigens.

### Targeting a neuron subtype

Finally, we asked if functionalized probes could be exploited to target different neuron subtypes. We modified probes with an antibody that binds to an extracellular region of dopamine receptor 2 (D2R), to distinguish D2R expressing neurons from the general population of neurons. Representative staining of D2R neurons in tissue sections with D2R antibody (D2R Ab)-modified (Fig. 4A) and unmodified mesh probes (Fig. 4B) near the CA1 region showed an increased association of D2R neurons with the D2R Ab modified probes that also extends into the sub-CA1 regions without endogenous D2R neurons (Fig. 4A-B, yellowed dashed boxes). The small number of D2R neurons detected in the sub-CA1 region of the D2R Ab-modified probes is consistent with trapping of D2R-expressing newborn neurons migrating from the dentate gyrus (*38*). The specific targeting of D2R neurons was confirmed by plotting the normalized D2R neuron density near the probe surface in the CA1 region (Fig. 4C, fig. S22).

**Fig. 4.**
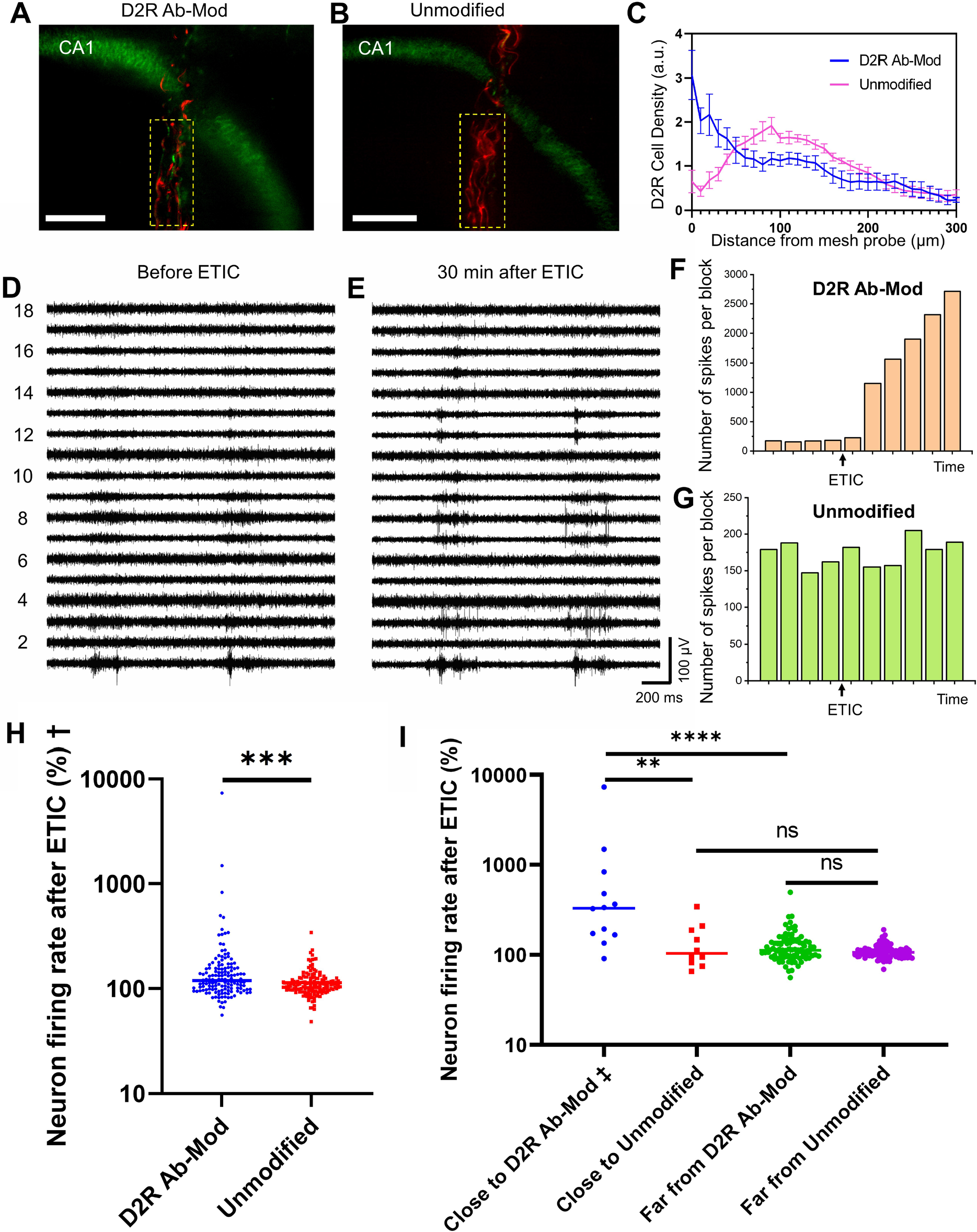
Targeting a neuron subtype. (**A**)-(**B**) Representative fluorescence images of D2R neurons adjacent to D2R Ab-modified (A, D2R Ab-Mod) and unmodified (B) mesh probes. The sub-CA1 regions without endogenous D2R neurons are highlighted in yellow dashed boxes. The D2R neurons are green and the mesh probe is red. Scale bars, 200 µm. (**C**) Normalized cell density distribution for D2R neurons as a function of distance from the probes (N = 6 mice for D2R Ab-modified and N = 5 mice for unmodified). Values are means ± s.e.m.. (**D**)-(**E**) Representative single-unit spike traces under anesthesia from a D2R Ab-modified probe immediately before and 30 min after ETIC injection. (**F**)-(**G**) Representative firing patterns of neurons recorded by D2R Ab-modified (F) and unmodified (G) probes. Black arrows denote the time of ETIC injection. Each bar represents spikes within a 5-min recording block. (**H**)-(**I**) Firing rate changes of neurons recorded by D2R Ab-modified and unmodified probes after ETIC injection (N = 3 mice). (H) all neurons, († 9 neurons recorded by D2R Ab probes started firing after ETIC not included), (I) neurons closely associated with probes (‡ 3 neurons near D2R Ab mesh probes started firing after ETIC not included) and neurons far away from probes. Values are medians, ns nonsignificant, ** P ≤ 0.01, *** P ≤ 0.001, **** P ≤ 0.0001, two-tailed, Mann-Whitney test.

Intracortical injection of 20 μg of D2R antagonist eticlopride (ETIC), which specifically increases the firing frequency of neurons with D2R (*39, 40*), was used to confirm the targeting of D2R neurons in electrophysiology recording. Injections were carried out ca. 1.5 mm away from the implanted mesh probes. The neural activity was recorded from mice under stable anesthesia for 20 min before and 30 min after ETIC injection to allow sufficient diffusion, from mesh probes with and without D2R Ab modification (N = 3 mice). Representative recordings from a D2R Ab mesh probe before and 30 min after ETIC injection (Fig. 4D-E) exhibited increased numbers of single-unit spikes post-ETIC injection, while the recordings from an unmodified mesh probe showed no clear difference before and after ETIC injection (fig. S23). Single neuron activity was tracked by counting the number of sorted spikes with principal component analysis from every channel as a function of time represented by 5-min recording blocks (fig. S24, fig. S25). Comparison of the representative changes of neurons recorded by a D2R Ab probe (Fig. 4F) and an unmodified probe (Fig. 4G) revealed that although the firing rate of the neurons recorded by the D2R Ab probe before ETIC injection remained constant, the firing rate after ETIC injection increased significantly and continued to increase for 30 min. The neurons recorded by the unmodified probe maintained a constant firing rate before and after ETIC injection. These changes for modified and unmodified probes are consistent with increased of firing of D2R neurons trapped at the modified probe surface and not increased firing of neurons that are simply connected in a circuit to remote D2R neurons.

To further examine the selective D2R neuron detection shown in the above experiments we analyzed all 151 neurons measured from three D2R Ab-modified probes and all 153 neurons measured from three unmodified probes, and plotted the firing rate changes by comparing the recording time block with the maximum number of spikes after ETIC injection to the one with the maximum number of spikes before ETIC injection (before as baseline at 100%) (Fig. 4H). The neurons recorded by the D2R Ab-modified probes showed a significantly higher increase in the firing rate. In addition, D2R Ab-modified probes recorded 9 neurons that were not firing initially but started firing after ETIC injection, which were excluded from the ratio calculation. As histology studies showed that D2R neurons preferentially clustered near the D2R Ab-modified probe surface, we confirmed this observation with electrophysiology by comparing the ratio of the firing rate changes between probe-adjacent neurons (with firing amplitudes greater than mean amplitude + s.d. (32.6 + 10.7 mV for neurons recorded by D2R Ab probes, 38.9 + 22.8 mV for unmodified probes) and probe-distant neurons (with amplitudes lower than the mean), and determined that neurons closely associated to the D2R Ab-modified probes showed significantly higher frequency increase (median: 332%) after ETIC injection than neurons near the unmodified probes (median: 103%) (Fig. 4I). The neurons located further away from the D2R Ab probes (median: 112%) did not show significant difference compared to those recorded by the unmodified probes (median: 106%). Taken together, these data demonstrate the ability to selectively target and closely associate neurons of a specific subtype to implanted electronics via binding of membrane surface antigens.

## Discussion

In this work, we have achieved cell type- and neuron subtype-specific targeting with mesh probes functionalized with specific peptides and antibodies. Chronic electrophysiological recording and time-dependent histology studies have demonstrated selective accumulation of the targeted cell types near the probe surface in a minimally invasive, nongenetic manner. Further development of these cell-targeting capabilities could open up many new opportunities in electrophysiology. First, specific modification of individual recording electrodes could trap single cells or subcellular structures for future *in vivo* studies. This methodology could also elucidate how the targeted cells integrate into local circuits. Second, incorporation of selective recording and stimulation electrodes (*12*) targeting neuron subtypes onto the mesh probe platform could help to decode local excitatory/inhibitory circuitries. Third, the versatility of our cell targeting strategies might be expanded by introducing artificial functional groups on the cell surface with bio-orthogonal chemistry (*41, 42*). We anticipate that achieving cell-type specific targeting and electrophysiology without requiring genetic modifications will expand and complement existing work based on genetically-modified organisms (*43, 44*).

## Materials and Methods

### Design and fabrication of mesh electronic probes

Mesh probes with double-sided electrodes and input/output (I/O) metal regions (fig. S1) were fabricated using standard photolithography as described previously (*28*). The key steps are as follows: (1) A 100-nm-thick nickel (Ni) sacrificial layer was thermally evaporated (Sharon Vacuum Co.) onto a 3-inch silicon (Si) wafer (n-type 0.005 Ωcm, 600 nm thermal oxide, NOVA Electronic Materials). (2) LOR 3A and S1805 (Microchem) were spin-coated at 4,000 rpm for 45 s and baked at 180 °C for 4 min and at 115 °C for 1 min, respectively. The photoresist was patterned by photolithography with a mask aligner (SUSS MA6 mask aligner, SUSS MicroTec) and developed (MF-CD-26, MicroChem Corp.) for 1 min. Following this photolithography process, a 100-nm-thick gold (Au) bottom I/O layer was deposited by thermal evaporation (Sharon Vacuum Co.), followed by a liftoff step (Remover PG, MicroChem). (3) The photolithography process in step (2) was repeated to define 100-nm-thick 20-μm-diameter platinum bottom electrode regions for E-beam evaporation (Denton Vacuum Co.), followed by a liftoff step. (4) For the bottom-passivation layer, negative photoresist SU-8 (SU-8 2000.5; MicroChem) labeled with Lissamine rhodamine B ethylenediamine (RhBen; ∼10 μg ml^-1^ (*14*); Thermo Fisher Scientific) was spin-coated on the Si wafer at 3,000 rpm for 30 s, pre-baked at 65 °C for 1 min and 95 °C for 1 min, and then patterned by photolithography with the mask aligner. After post-baking at 65 °C for 1 min and 95 °C for 1 min, The SU-8 resist was developed in SU-8 developer (Microchem), rinsed with isopropanol, dried, and hard baked at 185 °C for 1 h. (5) The photolithography process in step 2 was repeated to define metal interconnects, followed by 100-nm-thick Au deposition by thermal evaporation and liftoff. (6) The top passivation layer is then patterned using a similar procedure as in step 4, followed by hard baking at 195 °C for 1 h. The process in (3) was repeated to deposit the 100-nm thick top layer of platinum electrodes. To release mesh probes, the Si wafer was cleaned with oxygen plasma (50 W, 1 min) and immersed in a Ni etchant solution comprising 40 % FeCl_3_:39 % HCl:H_2_O = 1:1:10 on a 50 °C hot plate for 90 min. Released mesh probes were rinsed with deionized (DI) water three times and transferred to sterile 1× PBS solution (HyClone, GE Healthcare Life Sciences) before surface modification or implantation.

### Surface modification

The SU-8 polymer surface of mesh probes is modified at room temperature as follows. (1) A modification solution was made by dissolving 10 mM of 1-Ethyl-3-(3-dimethylaminopropyl)-carbodiimide (EDC, Sigma-Aldrich Inc.) and 75 mM of N-hydroxysuccinimide (NHS, Sigma-Aldrich Inc.) in 50 mM of 2-(N-morpholino)ethanesulfonic acid (MES, Sigma-Aldrich Inc.) buffer (pH = 6). (2) Mesh probes were transferred to PBS solution. A glass capillary tube with an inner diameter (ID) of 400 μm and outer diameter (OD) of 550 μm (Produstrial) is positioned near the I/O end of the mesh probe while it is suspended in solution. The syringe connected to the other end of the glass tube was manually retracted to draw in the mesh probe into the tube. The opening of the glass tube was gently tapped on Kimwipes (Kimtech Science) to remove the PBS solution while keeping the mesh probes attached to the inner wall. (4) The tube end closer to the mesh region was dipped into the modification solution, and once the solution covered the mesh region, the opposite end of the tube was covered with parafilm to prevent the modification solution from reaching the stem and I/O regions. (5) The mesh region was incubated in modification solution for 30 min, followed by rinsing in PBS 3 times. (6) 50 µM of peptide for targeting neurons (KKKAASIKVAV for histology and electrophysiology or KKKAASIKVAV-K(5FAM) for validation of surface functionalization, CPC Scientific Inc), 50 µM of antibodies for targeting microglia (anti-CD11b, ab184307, Abcam), or astrocytes (anti-EAAT2, AGC-022, Alomone Labs) or 5 µM of antibodies for targeting neurons with dopamine receptor 2 (D2R) (anti-D2DR, sc-5303, Santa Cruz Biotechnology) or without cell type specificity (rabbit anti-human IgG, 6145-01, Southern Biotech) was dissolved in PBS, steps (2)-were repeated to incubate the mesh region in antibodies or peptide. (7) Mesh probes were then transferred to 50 mM Tris buffer (pH = 8) for 30 min to quench the reaction and the probes were then stored in PBS at 4 °C. Modified probes were implanted within 1 week.

### *In vivo* brain implantation surgery

Adult (6–8 weeks, 20–25 g) male C57BL/6J mice (000664, The Jackson Laboratory) were used in the study. All procedures were approved by the Animal Care and Use Committee of Harvard University. The animal care and use programs at Harvard University meet the requirements of federal law (89–544 and 91–579) and NIH regulations and are also accredited by the American Association for Accreditation of Laboratory Animal Care (AAALAC). Before surgical procedures, animals were group-housed on a 12h:12h light:dark schedule in the Harvard University Biology Research Infrastructure (BRI) and fed with food and water ad libitum as appropriate. Animals were housed individually after surgical procedures.

*In vivo* implantation of the mesh probes into mouse brains was performed using a controlled stereotaxic injection method (*32*). All metal tools in direct contact with the animal subjects were bead-sterilized (Fine Science Tools) for 1 h before use, and all plastic tools in direct contact with the animal subjects were sterilized with 70 % ethanol and rinsed with sterile DI water and sterile PBS before use. The mesh probes were sterilized with 70 % ethanol followed by rinsing in sterile DI water and sterile PBS before implantation. Mice were anaesthetized by intraperitoneal injection of a mixture of 75 mg kg^-1^ of ketamine (Patterson Veterinary Supply) and 1 mg kg^-1^ dexdomitor (Orion). The degree of anesthesia was verified via toe pinch before surgery. A homeothermic blanket (Harvard Apparatus) was set to 37 °C and placed underneath the anaesthetized mouse. The anaesthetized mouse was placed in a stereotaxic frame (Lab Standard Stereotaxic Instrument, Stoelting Co.) equipped with two ear bars and one nose clamp. Puralube vet ointment (Dechra Pharmaceuticals) was applied to both eyes to prevent corneal damage. Hair removal lotion (Nair, Church & Dwight) was applied to the scalp for depilation and an alternating series of Betadine surgical scrub (Purdue Products) and 70 % alcohol was applied to sterilize the depilated scalp skin. A sterile scalpel was used to make a 5 mm longitudinal incision in the scalp along the sagittal sinus. The scalp skin was resected to expose a 5 mm×5 mm portion of the skull.

A 1-mm-diameter burr hole was made using a dental drill (Micromotor with On/Off Pedal 110/220, Grobet USA) according to the following stereotaxic coordinates: anteroposterior, 2 mm; mediolateral, 1.5 mm. A sterilized 0–80 set screw (McMaster-Carr Supply Company) was screwed into this burr hole to a depth of 500 μm to serve as the grounding and reference electrode. Metabond adhesive cement (Parkell) was applied to fix the screw/skull junction. Another 0.5-mm-diameter burr hole was made for the implantation of mesh probes according to the following stereotaxic coordinates: anteroposterior, -1.85 mm; mediolateral, 1.45 mm; dorsoventral, 2.50 mm. A flat, flexible cables (FFC, WM11484-ND, Digi-Key Electronics) with 32 conductors with a 0.50-mm pitch was attached with Metabond adhesive cement onto a head restrainer made with a 3D printer (Makerbot, Replicator 2x) using polylactic acid (PLA) filament. The head restrainer was attached and fixed on the skull with Metabond adhesive cement, the FFC near the cranium hole served as a substrate for I/O connection.

Mesh probes were implanted into the hippocampus using a controlled injection method. First, mesh probes were loaded into a sterile glass capillary tube with an ID of 300 μm and OD of 400 μm (Produstrial), which was inserted in a micropipette holder (1-HL-U, Molecular Devices LLC) fixed on a stereotaxic stage equipped with a motorized linear translation stage (860A motorizer and 460A linear stage, Newport) that could move the stereotaxic arm in the Z direction with a constant preset velocity ranging from 0.05 to 0.5 mms^-1^. The micropipette holder was connected to a 5 ml syringe which was pre-filled with PBS and mounted on a syringe pump (PHD 2000, Harvard Apparatus). The glass tube was inserted into the brain tissue to the targeted coordinates. Controlled injection was carried out by synchronizing the syringe pump with the motorized linear translation stage, with a PBS injection rate of 5–40 ml h^-1^ and a translational stage retraction velocity of 0.2–0.5 mm s^-1^. The volumetric flow rate and needle retraction velocity were adjusted such that the mesh region right above the burr hole, which was visualized through an eyepiece camera (DCC1240C, Thorlabs), remained stationary in the field of view. The total injection volume was 10–40 µl over the 2.5 mm length of injection. After implantation, the mesh probe is fixed at burr hole with Metabond adhesive cement. The 32 I/O pads were aligned with flowing DI water onto the 32 conductors on the FFC. After the I/O pads were dried, UV light glue (Visbella) was applied and cured with handheld UV flashlight (Vansky). For mesh probe implantation for histology studies without electrophysiology recording, the scalp was closed with 3M Vetbond tissue adhesive. Triple antibiotic ointment (Water-Jel Technologies) was applied copiously on the closure.

After surgery, each mouse was returned to a cage place on a 37 °C heating pad. The activity of the mouse was monitored every hour until it was fully recovered from anesthesia. Buprenex (Buprenorphine, Patterson Veterinary Supply) analgesia was given intraperitoneally at a dose of 0.05 mg per kg body weight every 12 h for up to 72 h post-surgery.

### Histology sample preparation and immunostaining

Mice with implanted mesh probes were anaesthetized and transcardially perfused with ice-cold 40 ml PBS and 40 ml 4% formaldehyde (Electron Microscopy Sciences) at specified times post-implantation, followed by decapitation. The scalp skin was removed, and the exposed skull was ground for 10–20 min at 10,000 r.p.m. using a high-speed rotary tool (Dremel). The brain was removed from the cranium and placed in 4% formaldehyde in PBS for 24 h and then transferred to PBS for at least 24 h. The brain was then embedded in 4% agarose (SeaPlaque agarose, Lonza) hydrogel, cut into blocks of 2 cm (length)×2 cm (width)×1 cm (height), then sectioned into 1-mm thick slices using a vibratome (VT1000S vibrating blade microtome, Leica). Fluorescence from the rhodamine within the mesh probes is visible through the tissue using a wide-field epifluorescence microscope (Olympus) when it is 50–100 μm from the tissue surface, which is used to adjust the sectioning such that one tissue slice includes the entire mesh probe.

After vibratome sectioning, brain slices with unmodified probes and probes modified with anti-EAAT2, anti-CD11b, IKVAV peptide, and control antibody were placed in PBS containing 4% (wt/vol) acrylamide (Sigma-Aldrich) and 0.25 % (wt/vol) VA-044 thermal polymerization initiator (Fisher Scientific) at 4 °C for 1 day. Brain slices with mesh probes modified with anti-D2R were placed in 1x PBS containing 4 % (wt/vol) acrylamide (Sigma-Aldrich) and 0.25 % (wt/vol) VA-044 thermal polymerization initiator (Fisher Scientific) and 0.5 % (wt/vol) paraformaldehyde at 4 °C for 1 day. The brain slices were placed in freshly made acrylamide/VA-044 solution immediately before placing the brain slices in X-CLARITY polymerization system (Logos Biosystems) for 3 h at 37 °C. After polymerization, any remaining gel from the tissue surface was removed and the slices were rinsed with PBST (PBS with 0.2 % Triton X-100, Thermo Fisher Scientific) before placing them in electrophoretic tissue clearing solution (Logos Biosystems) at 37 °C for 3–5 days until the samples were translucent. After clearing, the brain slices were placed in fresh PBST and gently shaken for 5h, with the PBST replaced by fresh solution every hour. The brain slices were incubated with 1:100–1:200 primary antibodies, rabbit anti-NeuN (ab177487, Abcam) and/or rat anti-GFAP (13-0300, Thermo Fisher Scientific) and/or goat anti-Iba1 (ab5076, Abcam) and/or mouse anti-D2R conjugated to Alexa-Fluor 488 (sc-5303 AF488, Santa Cruz Biotechnology) in PBST for 4 days at 4 °C with gentle shaking. After incubation, slices were placed in fresh PBST at 4 °C to let excess antibody diffuse out of the tissue. The PBST was replaced with fresh solution every 8 h over the course of 2 days. The samples were then incubated with 1:100–1:200 secondary antibodies, goat anti-rabbit Alexa Fluor 488 (ab150077, Abcam) or goat anti-rabbit Alexa Fluor 647 (ab150079, Abcam), or donkey anti-rabbit Alexa Fluor 488 (ab18723, Abcam) and/or donkey anti-rat Alexa Fluor 647 (ab150155, Abcam) and/or donkey anti-goat Alexa Fluor 594 (ab150132, Abcam) in PBST for 4 days at 4 °C. Fluorophore-conjugated D2R primary antibody was not targeted with a secondary antibody.

Brain slices were then incubated in the refractive index matching solution of 80% glycerol:20% PBS (glycerol, Sigma-Aldrich) at least 24 h before microscopy imaging. At least 3 hours before imaging, tissue samples were glued at their edge to the bottom of 50-mm-diameter Petri dishes by Devcon 5 minute epoxy (ITW Polymers Adhesives) and then incubated in the same refractive index matching solution of 80 % glycerol:20 % PBS (glycerol, Sigma-Aldrich).

### Microscopy imaging of brain slices

Fluorescence images were acquired on a Zeiss LSM 880 confocal microscope (Carl Zeiss Microscopy) with a 20× objective (numerical aperture 1.0, free working distance 5.6 mm) or with a Zeiss Lightsheet.Z1 Microscope. Images of cleared and immunostained brain slices were acquired using 405, 488, 561 and/or 633 nm lasers as the excitation sources for Alexa Fluor 405, Alexa Fluor 488, RhBen/Alexa Fluor 594 and/or Alexa Fluor 647, respectively. The corresponding bandpass filters were 411–496, 500–553, 562–624, 624–660 and/ or 677–755 nm, respectively. Confocal images were acquired by taking a tile scan together with Z stacks, with a voxel size of 0.2–0.6 μm(x)×0.2–0.6 μm(y)×0.4–2.5 μm(z) and tile scan overlap of 10%. Lightsheet images were taken using a z-resolution of 4 μm.

### Image processing and quantification

Tile scans and Z-stacks of images were stitched using Zen Blue software (Zeiss). Images were then loaded into ImageJ (National Institute of Health), channels were split, and then saved into individual h5 files using Ilastik ImageJ plugin to export. The signal is then segmented in these images into individual objects by using Ilastik’s pixel classification and object classification workflow (*34, 35*). This involves manually assigning ground truth for pixels that are signal and for pixels that are not signal in individual slices of the image stack, and using these assignments to teach a classifier to distinguish fluorescent labeling from noise. We trained the classifier on each channel (Iba1, NeuN, GFAP, Dopamine receptor 2, and rhodamine-mesh) in a minimum of 1000 imaging planes across several independent imaging volumes, then applied the classifier with bulk processing to the rest of the data and generated probability masks that represent the likelihood that each pixel is signal. These probability masks were then used in the object classification workflow to separate the individual microglia, neurons, astrocytes, dopamine receptor 2 neurons, and mesh electronics implant, and then generate an information table about their center and size. Matlab (Mathworks) was then used to quantify and plot the shortest distance of all segmented cells from the mesh electronics. Imaris (Oxford Instruments) was used to draw a virtual surface around the CA1 (fig. S26), which was then exported as an OME tiff with pixel value 1 for the region within the surface. This CA1 tiff was then incorporated into the Matlab code to identify which segmented cells were within the CA1 of the hippocampus.

### Chronic electrophysiology recording

The electrophysiology of mice with implanted mesh probes was recorded for 10 min on days 1, 3, 5, 7, and 14 post-implantation, except for those used for D2R modulation. During recording, mice were restrained in a Tailveiner restrainer (Braintree Scientific) while their head restrainers were fixed and the head-mounted FFCs were connected to an Intan RHD 2132 amplifier evaluation system (Intan Technologies) through a homemade printed circuit board. 0–80 set screws implanted through the skull were used as reference electrodes as described previously (*12*). Electrophysiological recordings were acquired with a 20 kHz sampling rate and a 60 Hz notch filter. Matlab (Mathworks) was used to plot and sort the spikes (*12*). Statistical analyses were performed using GraphPad Prism 9.

### Modulation of D2R neuron activity

21-27 days post-implantation, mice were anaesthetized by intraperitoneal injection of a mixture of 75 mg kg^-1^ of ketamine (Patterson Veterinary Supply) and 1 mg kg^-1^ dexdomitor (Orion). The degree of anesthesia was verified via toe pinch before surgery. Puralube vet ointment (Dechra Pharmaceuticals) was applied on both eyes to prevent corneal damage. A Deltaphase isothermal pad (Braintree Scientific Inc) was heated to 39 °C and placed underneath the anaesthetized mouse to maintain the mouse body temperature at 37 °C throughout surgery and electrophysiology recording. An alternating series of Betadine surgical scrub (Purdue Products) and 70% alcohol was applied to sterilize the skull.

A burr hole was made for intracortical drug administration according to the following stereotaxic coordinates: anteroposterior, -0.35 mm; mediolateral, 1.45 mm (1.5 mm away from the implanted mesh probes). Drug solutions were made by dissolving D2R antagonist eticlopride (2 mg in 1 ml, Sigma-Aldrich) in sterile saline. 10 μL of the drug solutions were injected 1-mm deep into the burr hole using 27 G needle. Before and after drug administration, neuronal activity was recorded for 20 and 30 min, respectively. After recording, the burr hole was closed by Metabond adhesive cement (Parkell), and the mouse was returned to a cage place on a 37 °C heating pad. The activity of the mouse was monitored every hour until it was fully recovered from anesthesia. Buprenex (Buprenorphine, Patterson Veterinary Supply) analgesia was given intraperitoneally at a dose of 0.05 mg per kg body weight every 12 h for up to 72 h post-surgery.

## Acknowledgments

This work was performed in part at the Harvard University Center for Nanoscale Systems (CNS), a member of the National Nanotechnology Coordinated Infrastructure Network (NNCI), which is supported by the National Science Foundation under NSF award no. ECCS-2025158. We thank the Harvard Center for Biological Imaging (RRID:SCR_018673) for infrastructure and support. This work was supported by Air Force Office of Scientific Research FA9550-18-1-0469 (C.M.L), Lieber Research Group (C.M.L.), American Heart Association 23POST1018301 (A.Z.), National Institutes of Health K99AG068602 (T.J.Z.), and Harrison Gardner, Jr. Award (T.J.Z.).

## Author contributions

Conceptualization: AZ, TJZ, CML

Methodology: AZ, TJZ, CML

Investigation: AZ, TJZ, CML

Visualization: AZ, TJZ

Funding acquisition: CML, AZ, TJZ

Project administration: CML

Supervision: CML

Writing – original draft: AZ, TJZ, CML

Writing – review & editing: AZ, TJZ, CML

## Supplementary Materials

Figs. S1 to S26

