## Supplementary Information for "Biochemically-functionalized probes for cell type-specific targeting and recording in the brain"

This file contains:

Supplementary Figures 1 – 26

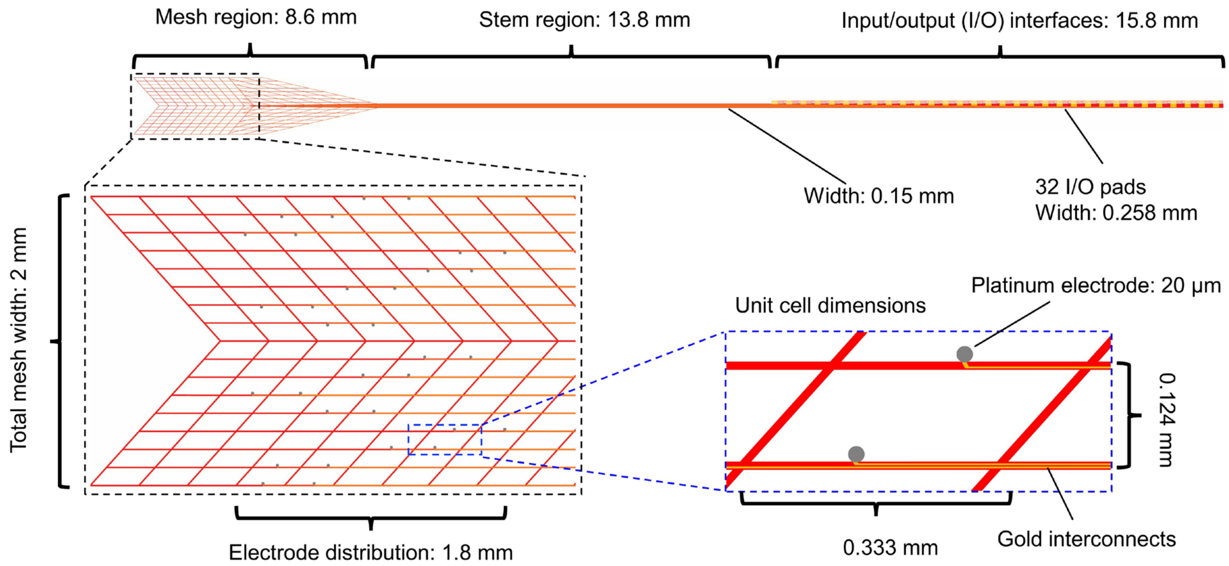


**Fig. S1.**

**Structure of the mesh probe.** Top, schematic of a mesh probe with the ultra-flexible mesh device region at left, the flexible stem in the middle, and double-sided input/output (I/O) regions at right. Bottom, magnified views of the 32 double-sided platinum recording electrodes (left) and one unit cell (right) in the mesh region. The mesh device region is partially delivered by syringe injection into the hippocampus, while the I/O interfaces are fixed outside on the skull, providing an electrical interface to the external recording instrument. SU-8 ribbons and stem, red; platinum electrodes, grey; gold interconnects, gold.

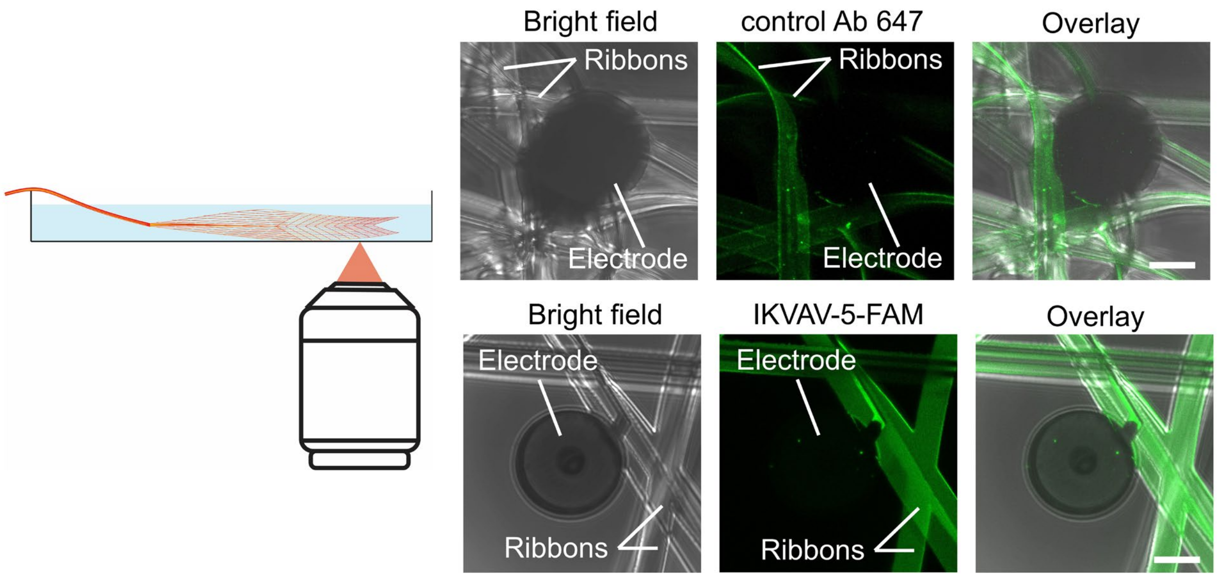


**Fig. S2.**

**Characterization of the modification layers on the SU-8 ribbons.** Left, schematic of the confocal imaging of modified mesh probes suspended in PBS. Right, mesh probes modified by control Ab, rabbit anti-human IgG, and then labeled with secondary antibody with Alexa 647 fluorophores (top) and IKVAV peptide labeled with 5-FAM fluorophores (bottom) using EDC/NHS covalent coupling chemistry (see materials and methods). Bright field images show SU-8 ribbons and platinum electrodes, while the fluorescence images show conjugated fluorophores are uniformly distributed on the modified SU-8 ribbons for both the antibody and peptide modifications. These results are representative as well of the antibodies used to target astrocytes (anti-EAAT2) and microglia (anti-CD11b). Scale bars, 10 µm.

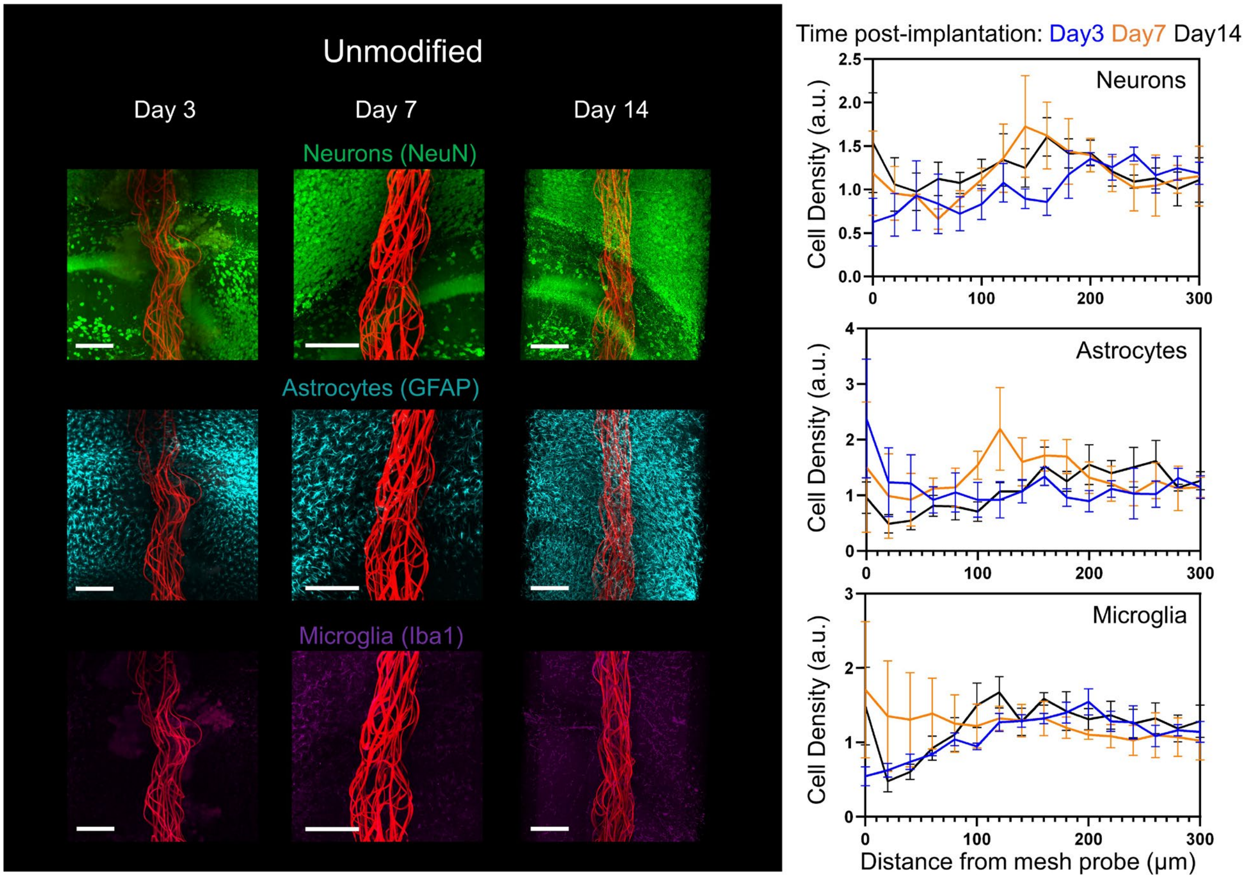


**Fig. S3.**

**Time-dependent 3D histology studies of unmodified mesh probes.** Left, time-dependent histology studies 3, 7, and 14 days post-implantation of NeuN labelling of neurons (top), GFAP labelling of astrocytes (middle), and Iba1 labelling of microglia (bottom) distribution adjacent to unmodified mesh probes. Scale bars, 200 μm. Right, normalized cell density for neurons (top), astrocytes (middle), and microglia (bottom) as a function of distance from the mesh probes on days 3 (blue), 7 (orange), and 14 (black) post-implantation (at least N = 3 independent mice at each time point). Values are means ± s.e.m.. The unmodified mesh probes did not show significant influence on the distribution of neurons, astrocytes, or microglia.

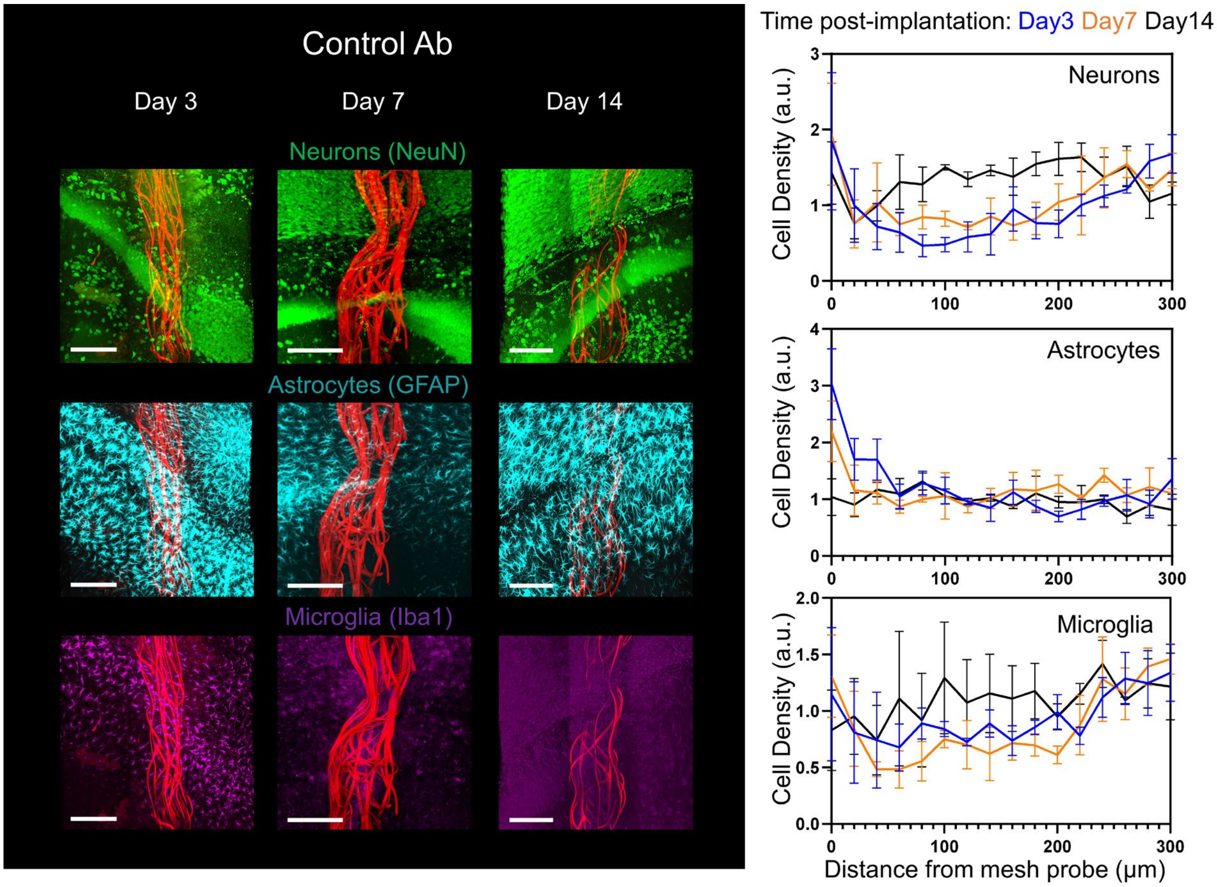


**Fig. S4.**

**Time-dependent 3D histology studies of mesh probes modified with control antibody.** Left, time-dependent histology studies 3, 7, and 14 days post-implantation of NeuN labelling of neurons (top), GFAP labelling of astrocytes (middle), and Iba1 labelling of microglia (bottom) distribution adjacent to mesh probes modified with control antibody. Scale bars, 200 µm. Right, normalized cell density for neurons (top), astrocytes (middle), and microglia (bottom) as a function of distance from the mesh probes on days 3 (blue), 7 (orange), and 14 (black) post-implantation (N = 3 independent mice at each time point). Values are means ± s.e.m.. The control Ab-modified mesh probes did not have a significant influence on the distributions of neurons, astrocytes, or microglia.

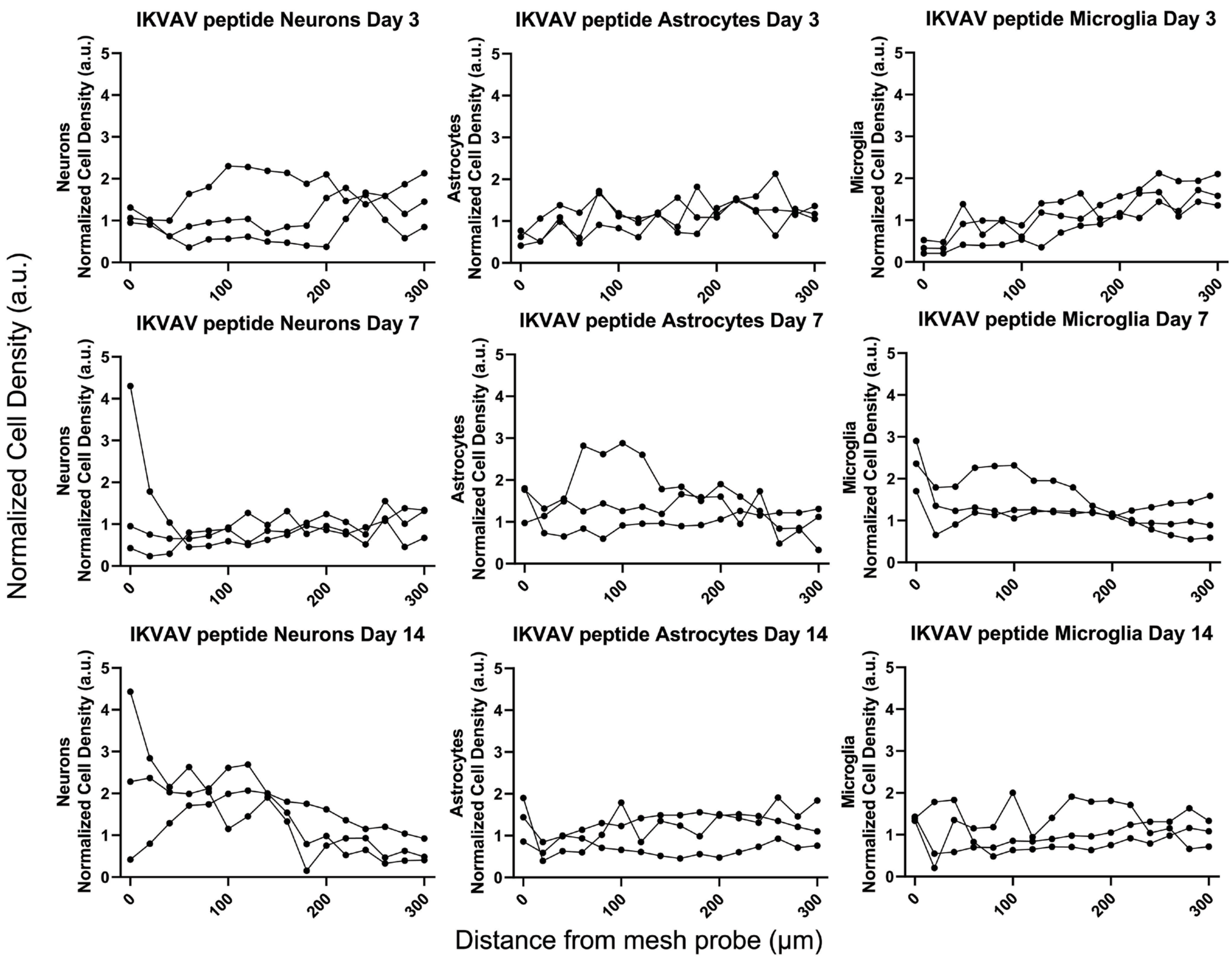


**Fig. S5.**

**Cell density distribution plots for individual samples in Fig. 2D.** Normalized cell density for neurons (left), astrocytes (middle), and microglia (right) in the CA1 region as a function of distance from surface of mesh probes modified with IKVAV peptide, on days 3 (left), 7 (middle), and 14 (right). The density of astrocytes and microglia always remained consistent with the baseline, while the neuron density increased 14 days post-implantation.

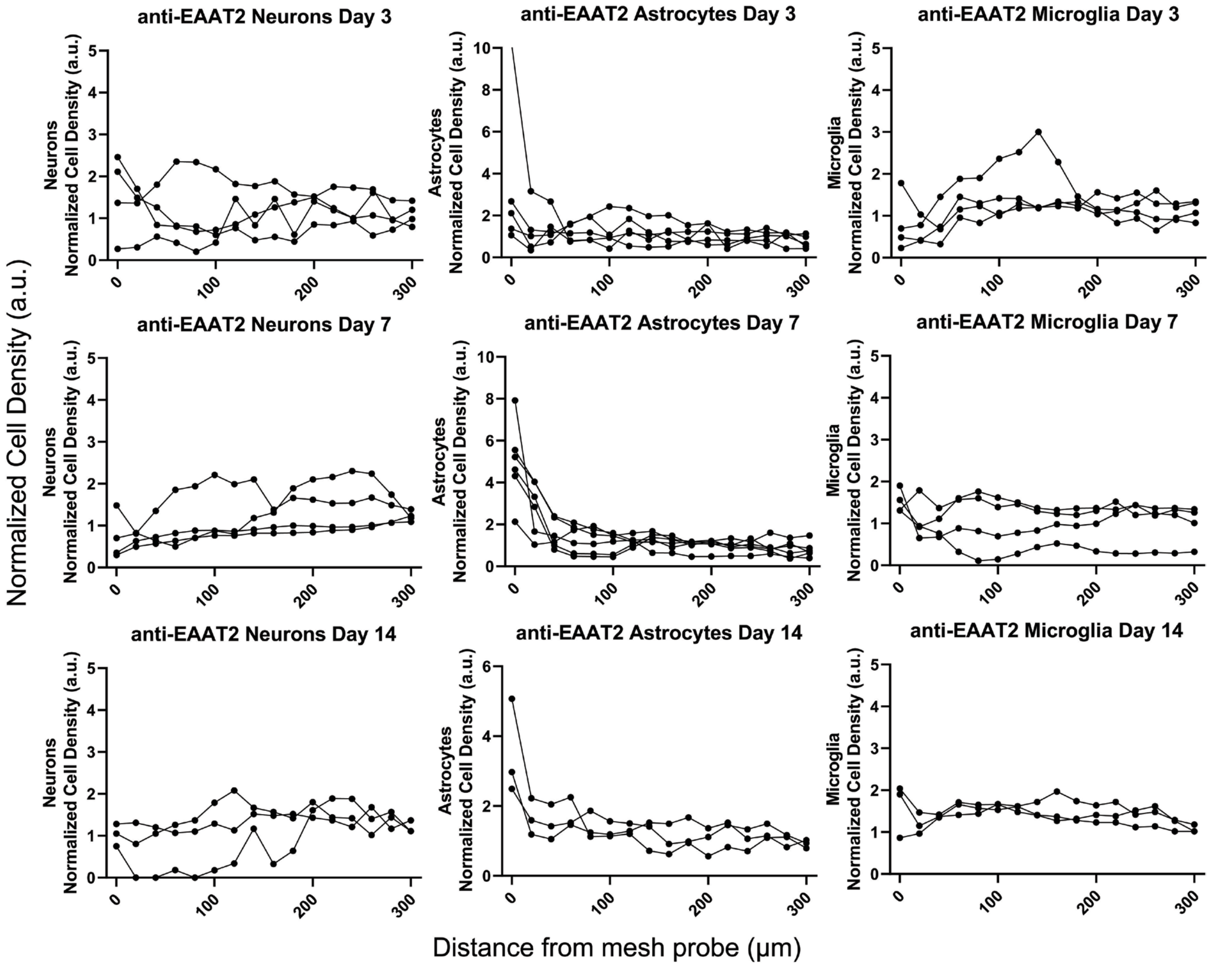


**Fig. S6.**

**Cell density distribution plots for individual samples in Fig. 2E.** Normalized cell density for neurons (left), astrocytes (middle), and microglia (right) in the CA1 region as a function of distance from surface of mesh probes modified with anti-EAAT2, on days 3 (left), 7 (middle), and 14 (right). An increased association of astrocytes was observed at all time points from day 3 to day 14, the microglia density remained close to the baseline level.

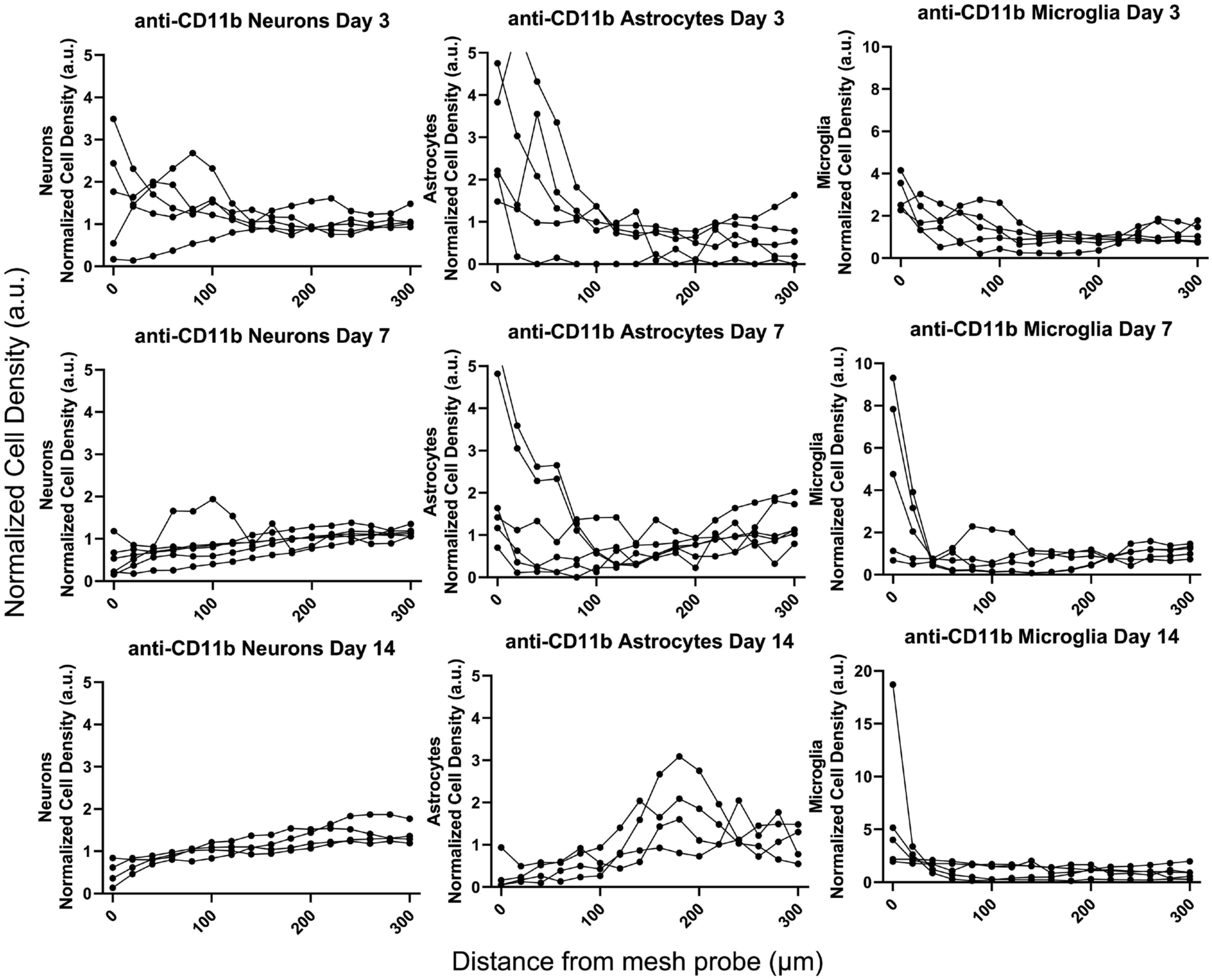


**Fig. S7.**

**Cell density distribution plots for individual samples in Fig. 2F.** Normalized cell density for neurons (left), astrocytes (middle), and microglia (right) in the CA1 region as a function of distance from surface of mesh probes modified with anti-CD11b, on days 3 (left), 7 (middle), and 14 (right). An increased association of microglia was observed at all time points from day 3 to day 14, while the astrocyte density exhibited an increase at early times and dropped below the baseline by day 14.

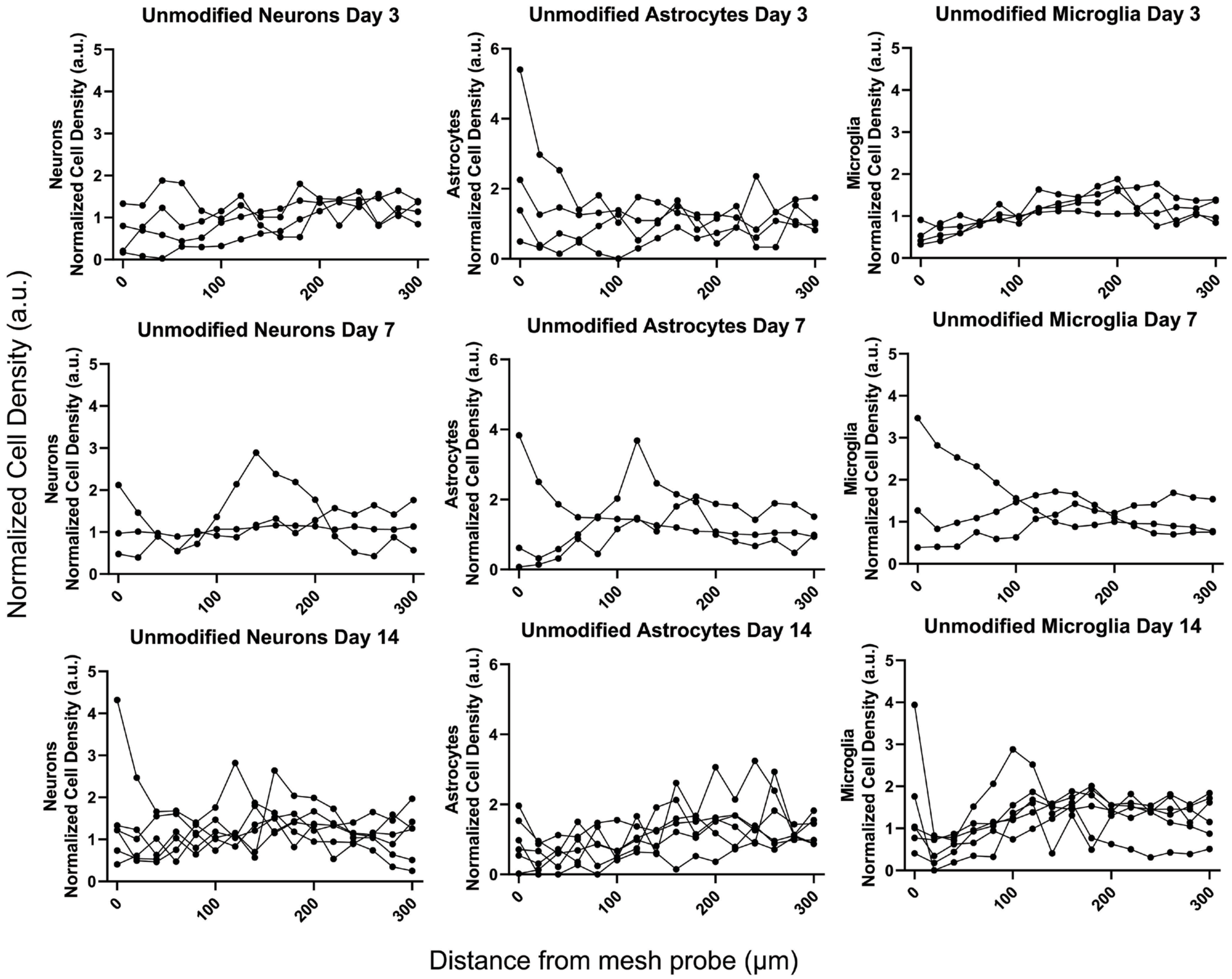


**Fig. S8.**

**Cell density distribution plots for individual samples in fig. S3.** Normalized cell density for neurons (left), astrocytes (middle), and microglia (right) in the CA1 region as a function of distance from surface of unmodified mesh probes, on days 3 (left), 7 (middle), and 14 (right). No significant influence on the distribution of neurons, astrocytes, or microglia was observed.

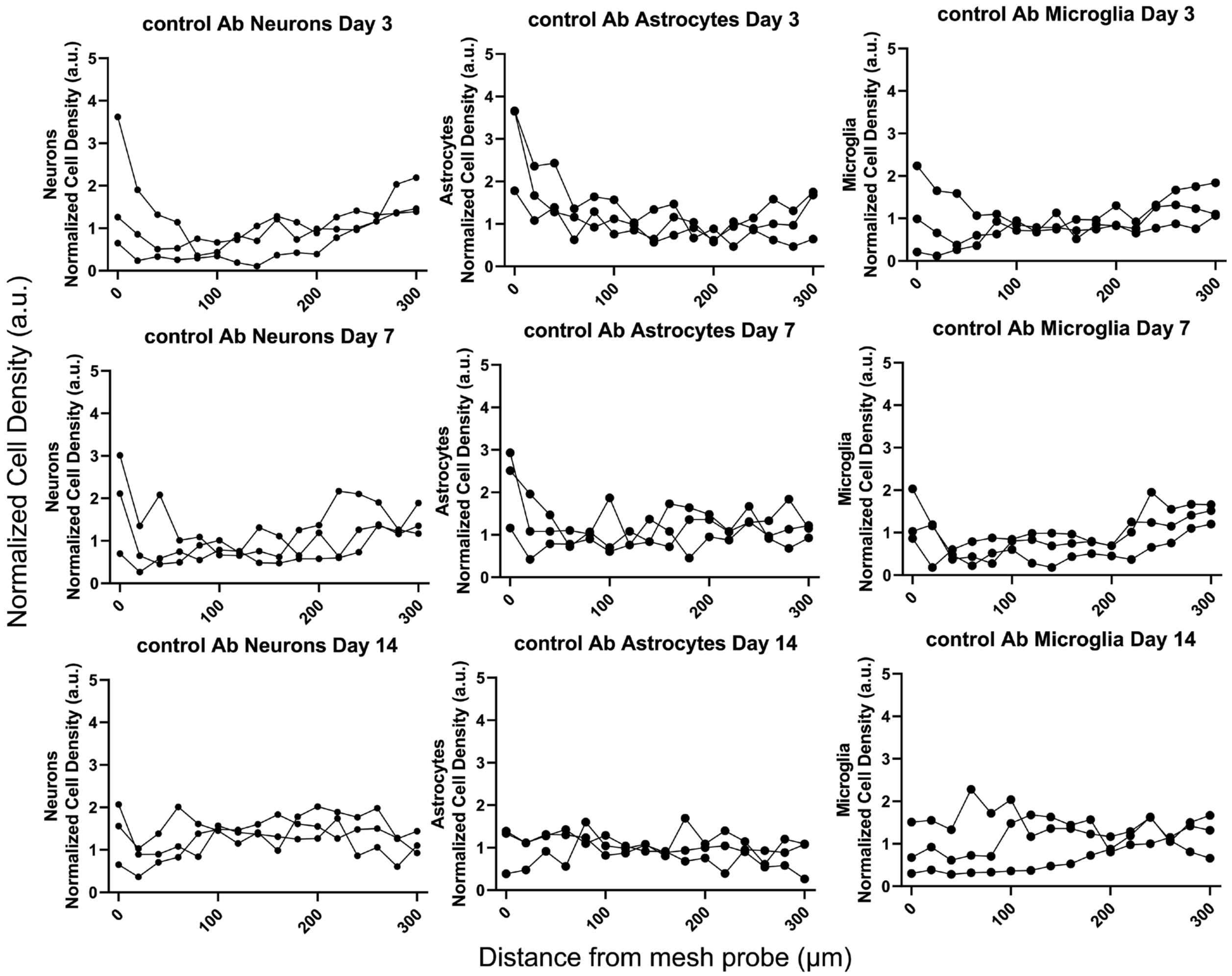


**Fig. S9.**

**Cell density distribution plots for individual samples in fig. S4.** Normalized cell density for neurons (left), astrocytes (middle), and microglia (right) in the CA1 region as a function of distance from surface of mesh probes modified with control antibody, on days 3 (left), 7 (middle), and 14 (right). No significant influence on the distribution of neurons, astrocytes, or microglia was observed.

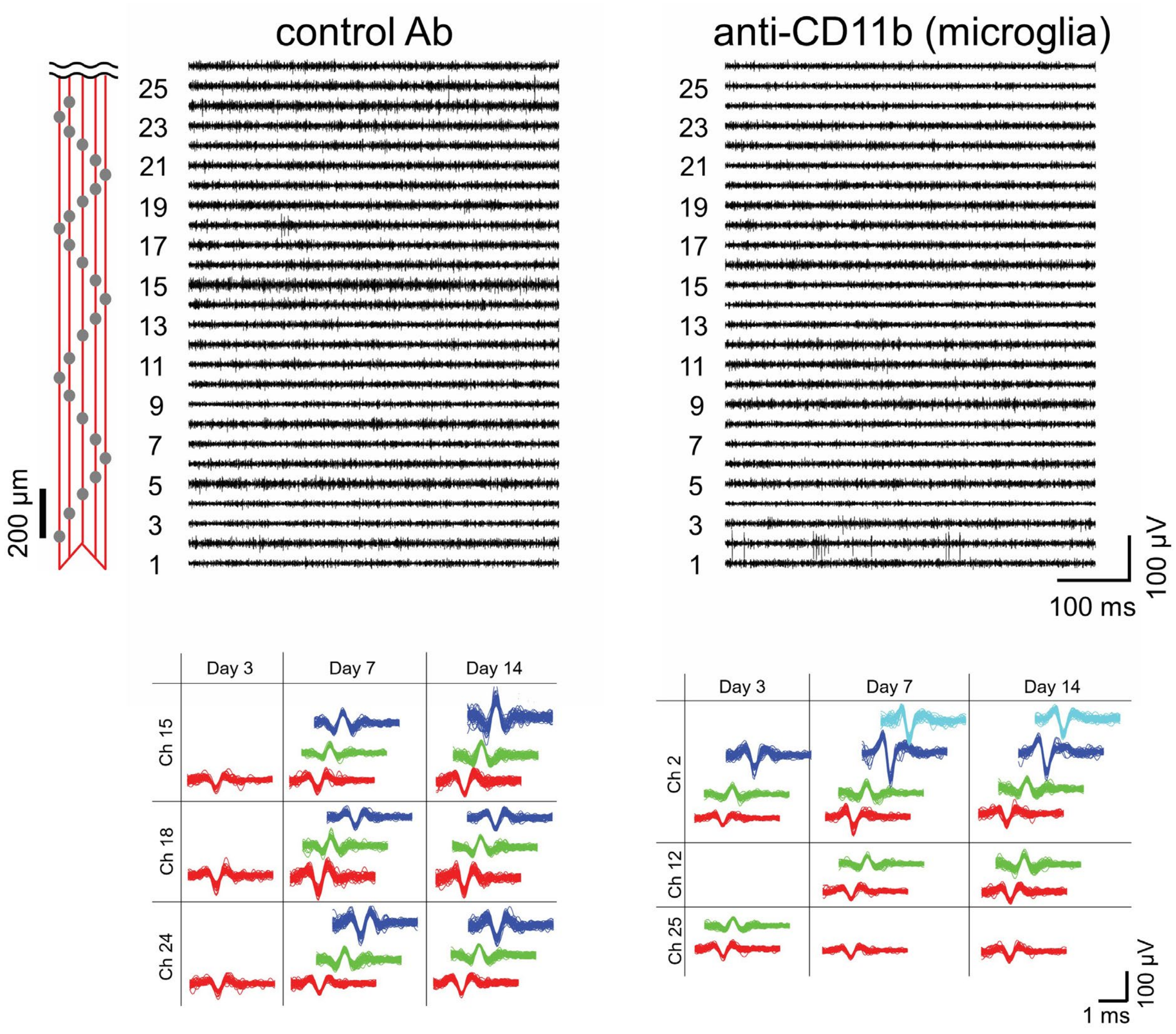


**Fig. S10.**

**Electrophysiological recording from mesh probes modified with control Ab without targeting specificity and anti-CD11b targeting microglia.** Top, representative 26-channel single-unit spike traces 14 days post-implantation from mesh probes modified with control antibody without specific targeting (left) and anti-CD11b for targeting microglia (right). Recordings were made from head-fixed, awake mice. The x and y axes represent recording time and voltage. The relative positions of the recording electrodes are marked by grey dots in the schematic (leftmost panel). Channels with smaller numbers are implanted deeper. Bottom, time evolution principal component analysis clustered single unit spikes from three representative channels. For each channel, each distinct color in the sorted spikes represents a unique identifiable neuron. The control Ab-modified probes showed a similar level of spontaneous firing as that recorded by the unmodified probes, while the anti-CD11b-modified probes showed a decreased level of spontaneous firing.

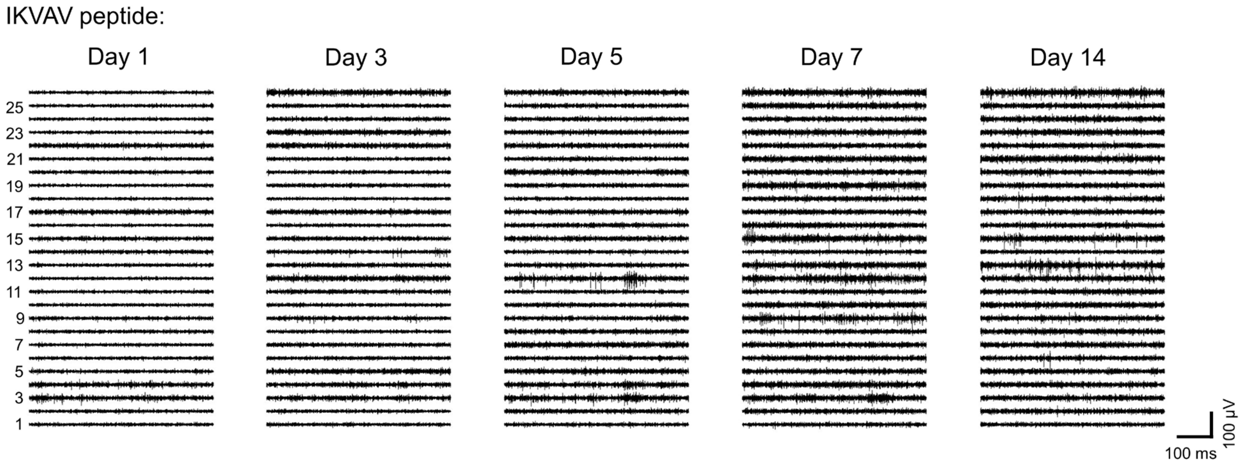


**Fig. S11.**

**Time-dependent recording from a mesh probe modified with IKVAV peptide.**
Representative 26-channel single-unit spike traces from the mesh probe with IKVAV peptide modification shown in Fig. 3A on days 1, 3, 5, 7, 14 post-implantation. The x and y axes represent recording time and voltage. Channels with smaller numbers are implanted deeper. The recorded spontaneous activity increased over time.

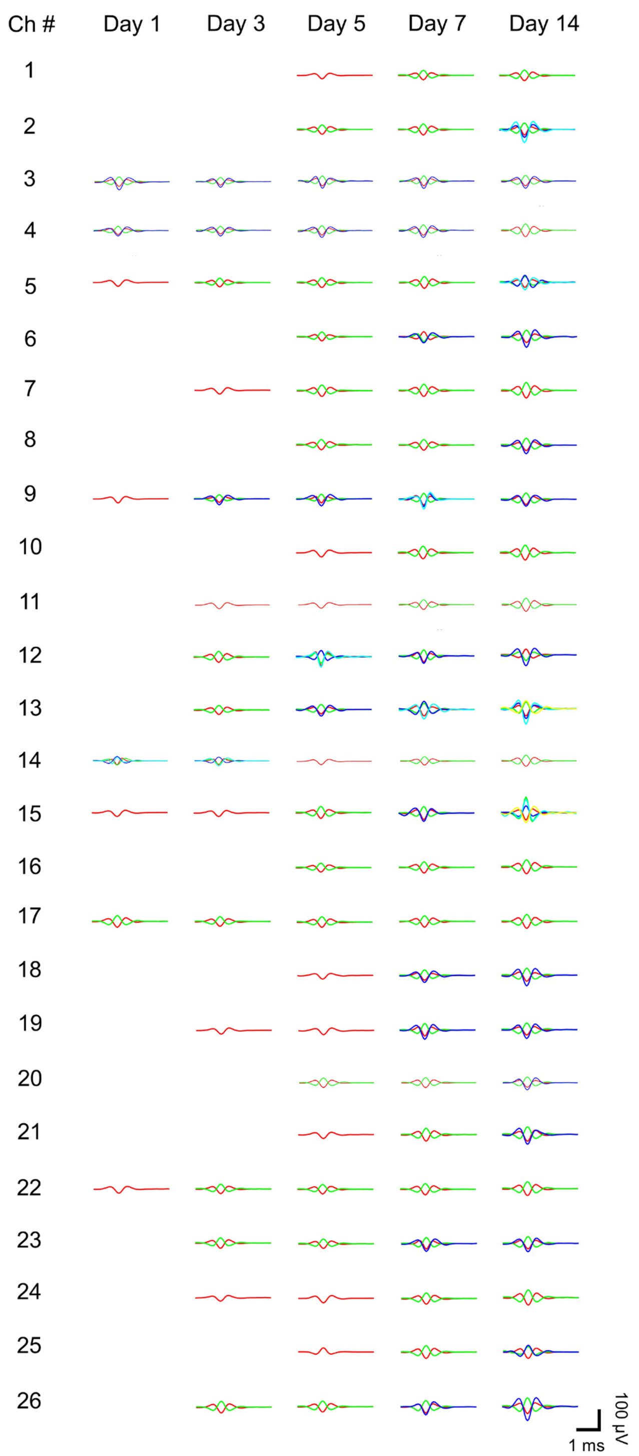


**Fig. S12.**

**Clustered single unit activity from all channels in fig. S11.** Time evolution of average single unit spike waveforms clustered by principal component analysis. For each channel, each distinct color in the sorted spikes represents a unique identifiable neuron.

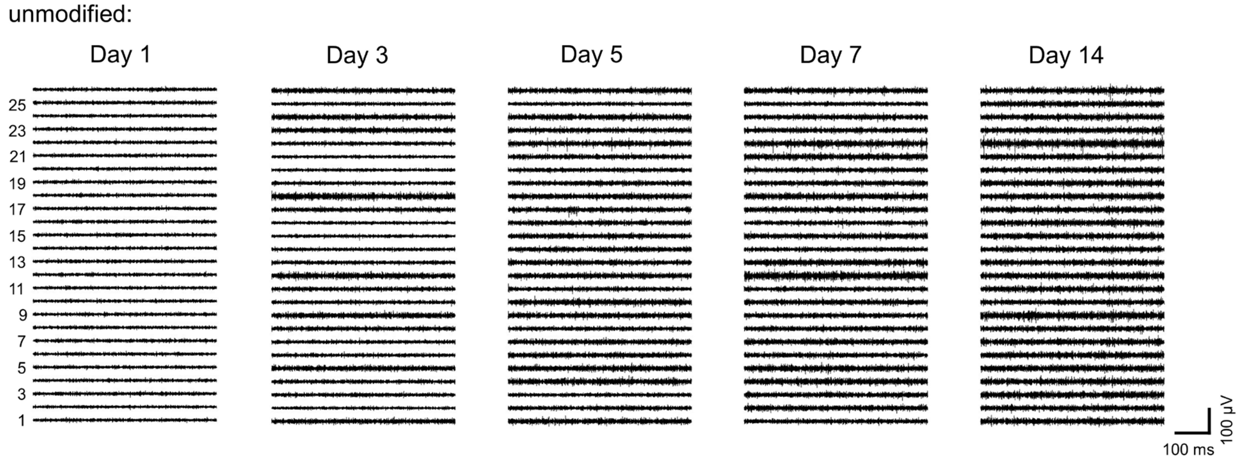


**Fig. S13.**

**Time-dependent recording from an unmodified mesh probe.** Representative 26-channel single-unit spike traces from the unmodified mesh probe shown in Fig. 3B on days 1, 3, 5, 7, 14 post-implantation. The x and y axes represent recording time and voltage. Channels with smaller numbers are implanted deeper. The recorded spontaneous activity increased over time.

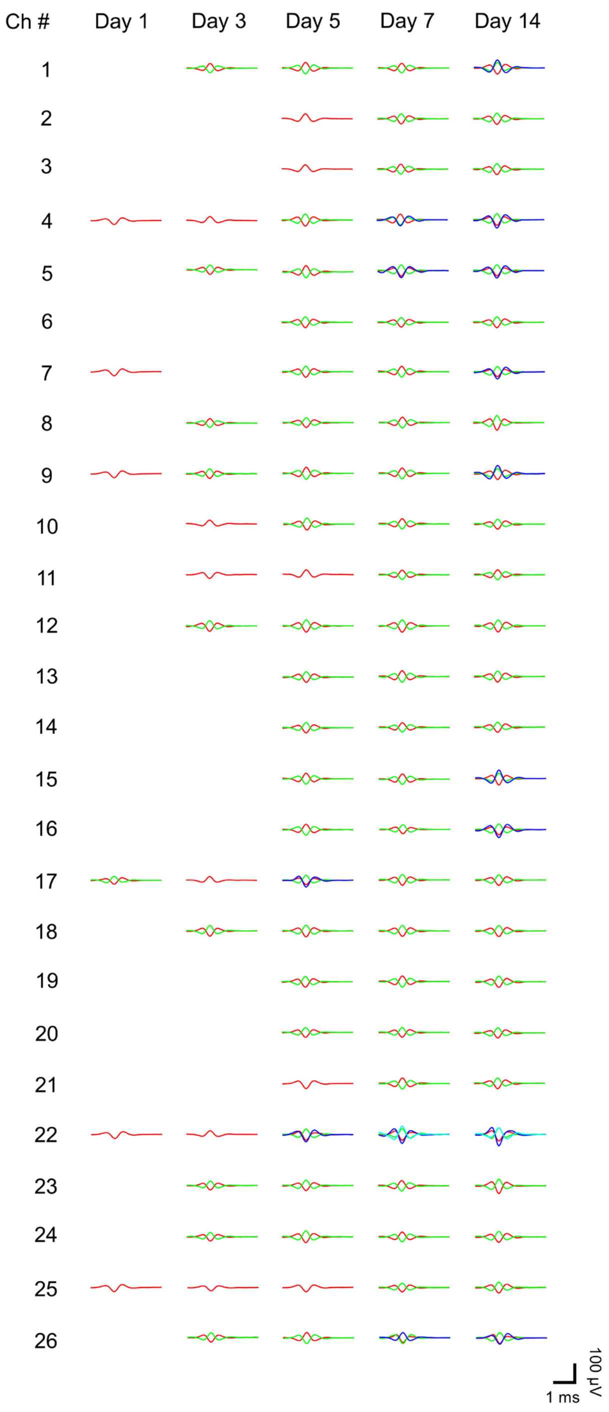


**Fig. S14.**

**Clustered single unit activity from all channels in fig. S13.** Time evolution of average single unit spike waveforms clustered by principal component analysis. For each channel, each distinct color in the sorted spikes represents a unique identifiable neuron.

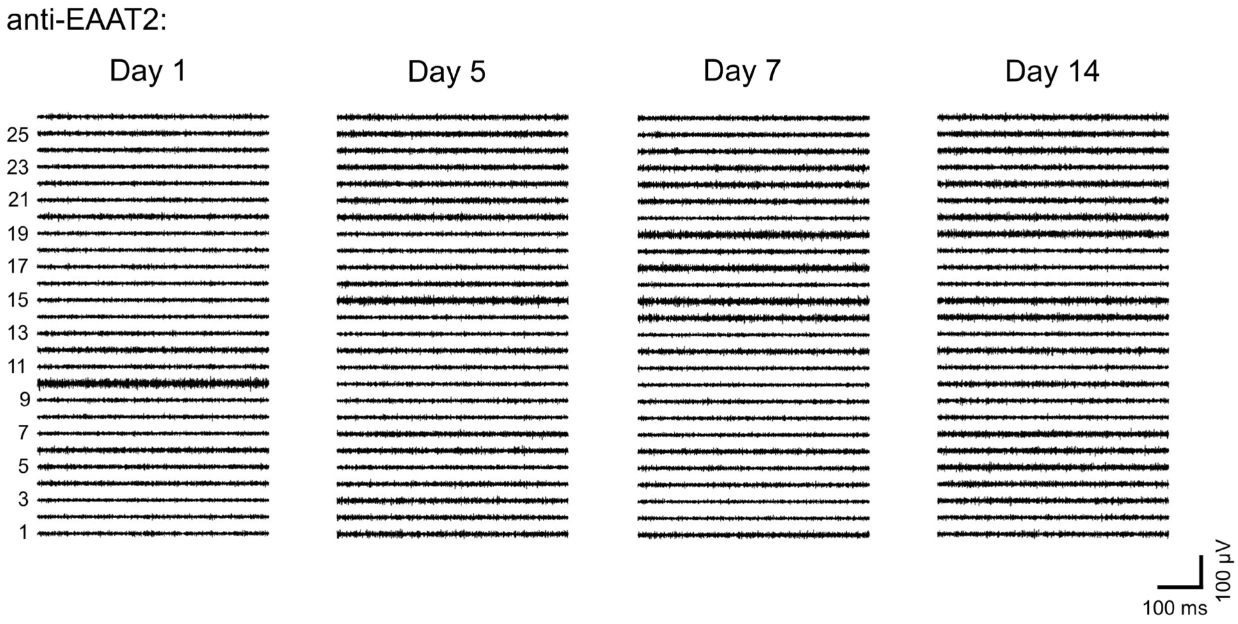


**Fig. S15.**

**Time-dependent recording from a mesh probe modified with anti-EAAT2.** Representative 26-channel single-unit spike traces from the mesh probe with anti-EAAT2 modification shown in Fig. 3C on days 1, 5, 7, 14 post-implantation. The x and y axes represent recording time and voltage. Channels with smaller numbers are implanted deeper. The recorded spontaneous activity did not show significant increase over time.

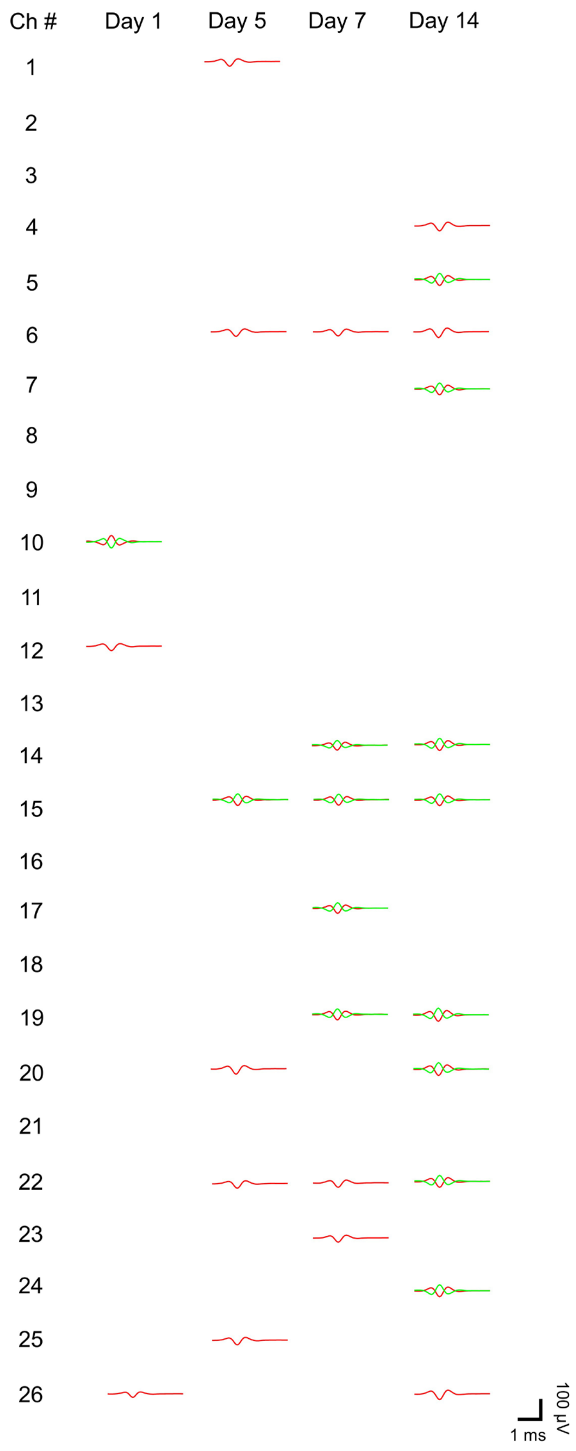


**Fig. S16.**

**Clustered single unit activity from all channels in fig. S15.** Time evolution of average single unit spike waveforms clustered by principal component analysis. For each channel, each distinct color in the sorted spikes represents a unique identifiable neuron.

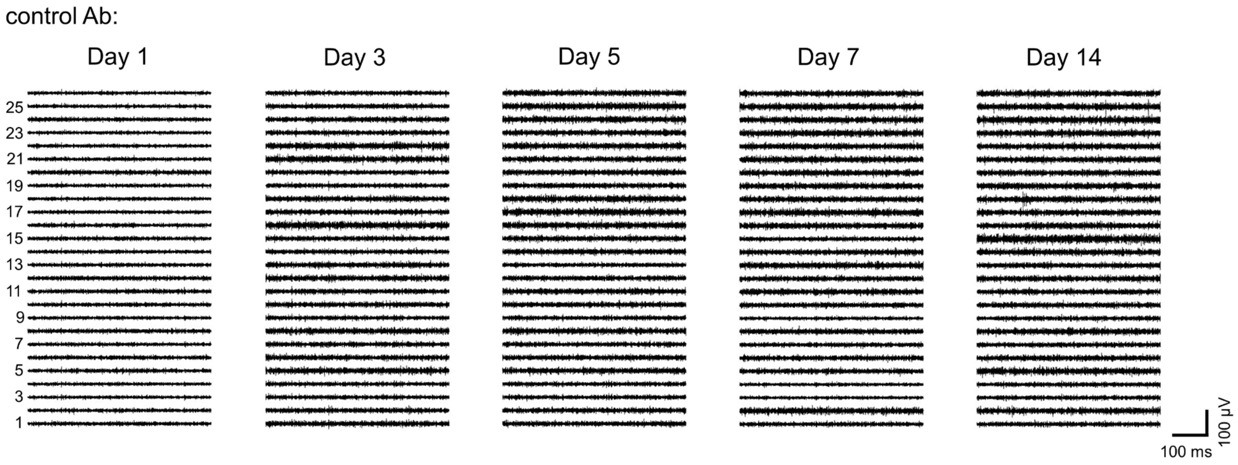


**Fig. S17.**

**Time-dependent recording from a mesh probe modified with control antibody.** Representative 26-channel single-unit spike traces from the mesh probe with control antibody modification shown in fig. S10 on days 1, 3, 5, 7, 14 post-implantation. The x and y axes represent recording time and voltage. Channels with smaller numbers are implanted deeper. The recorded spontaneous activity increased over time.

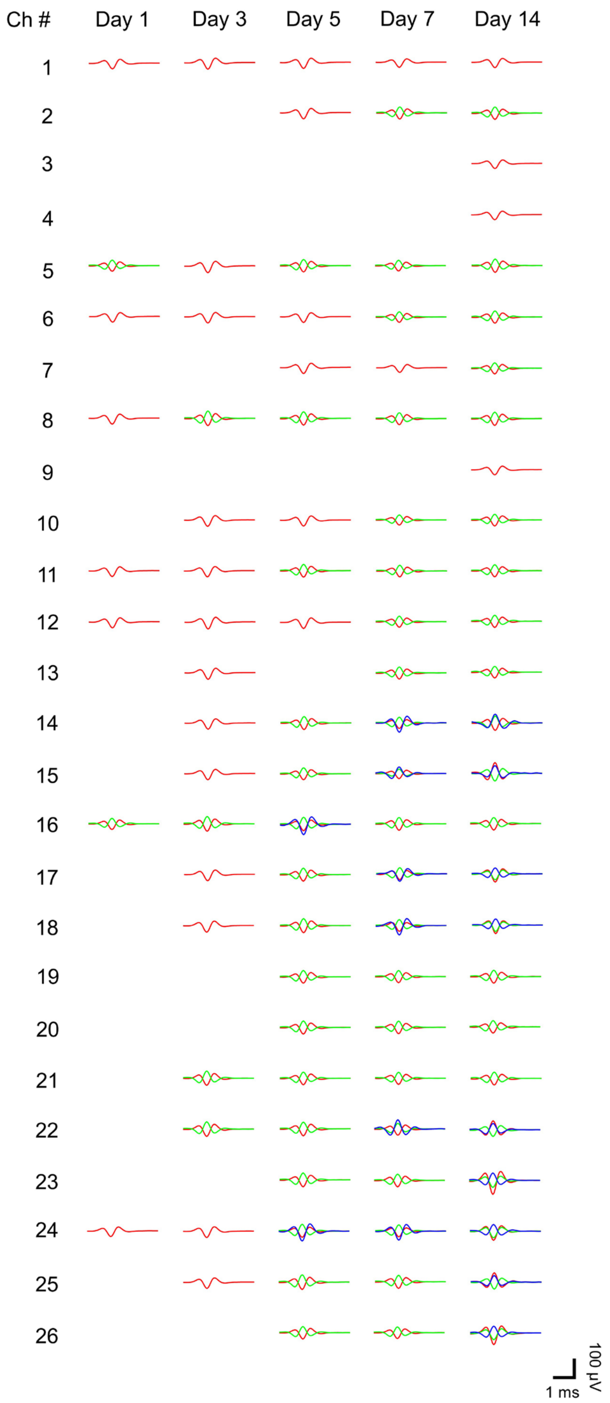


**Fig. S18.**

**Clustered single unit activity from all channels in fig. S17.** Time evolution of average single unit spike waveforms clustered by principal component analysis. For each channel, each distinct color in the sorted spikes represents a unique identifiable neuron.

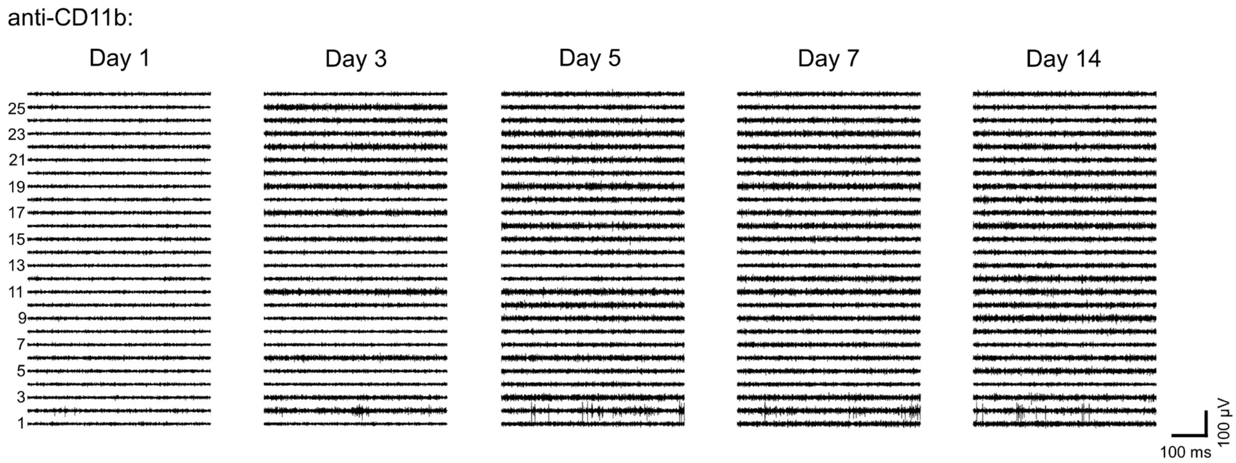


**Fig. S19.**

**Time-dependent recording from a mesh probe modified with anti-CD11b.** Representative 26-channel single-unit spike traces from the mesh probe with anti-CD11b modification shown in fig. S10 on days 1, 3, 5, 7, 14 post-implantation. The x and y axes represent recording time and voltage. Channels with smaller numbers are implanted deeper. The recorded spontaneous activity increased over time, although slower than that recorded by unmodified probes.

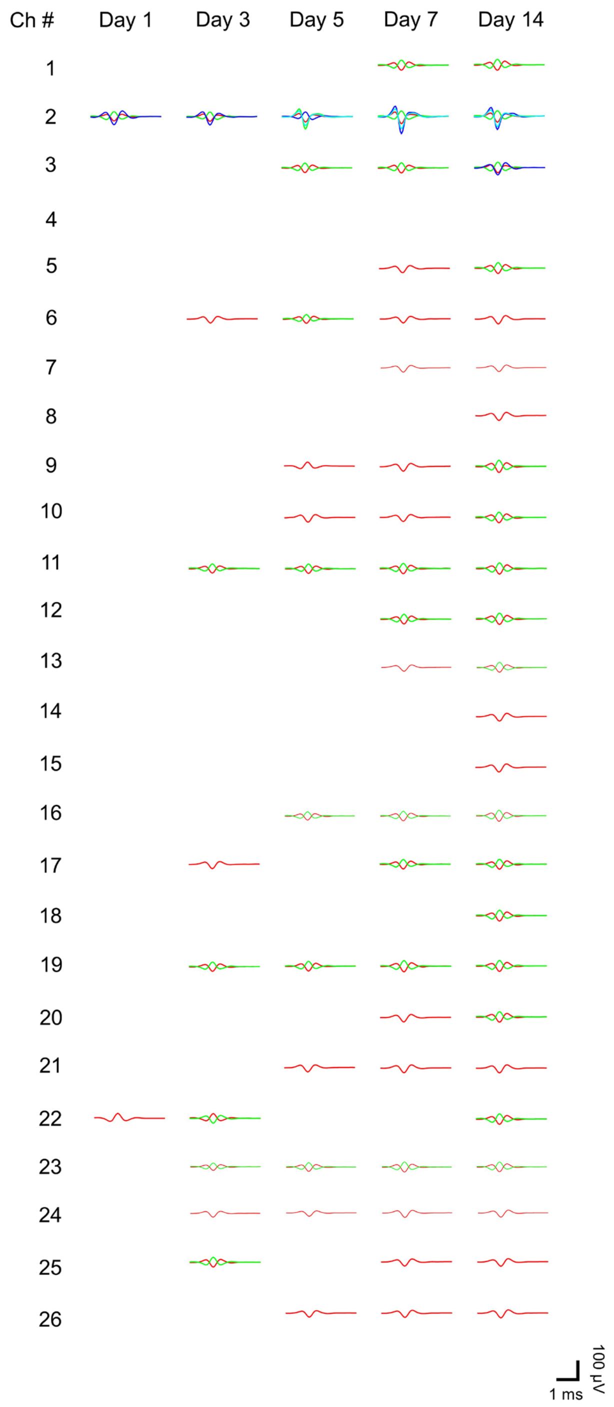


**Fig. S20.**

**Clustered single unit activity from all channels in fig. S19.** Time evolution of average single unit spike waveforms clustered by principal component analysis. For each channel, each distinct color in the sorted spikes represents a unique identifiable neuron.

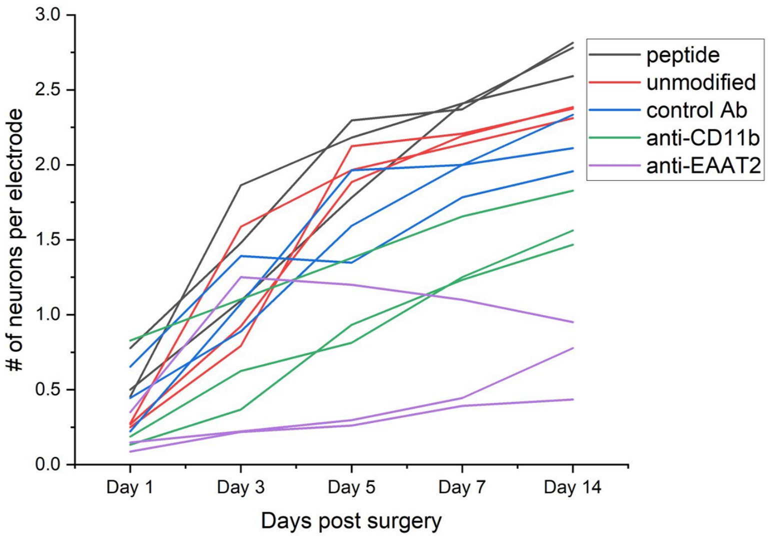


**Fig. S21.**

**Number of neurons recorded per electrode vs. days post-implantation.** The average number of distinct neurons recorded per electrode from day 1 to day 14 post-implantation from mesh probes with different modification. Each line represents one mesh probe. Data on day 14 was plotted in Fig. 3D. The number of neurons recorded per electrode increased over time for IKVAV peptide-modified, unmodified, and control Ab-modified probes. The number also increased slightly for anti-CD11b-modified probes. The anti-EAAT2 showed the lowest number of recorded neurons and the slowest increase over time.

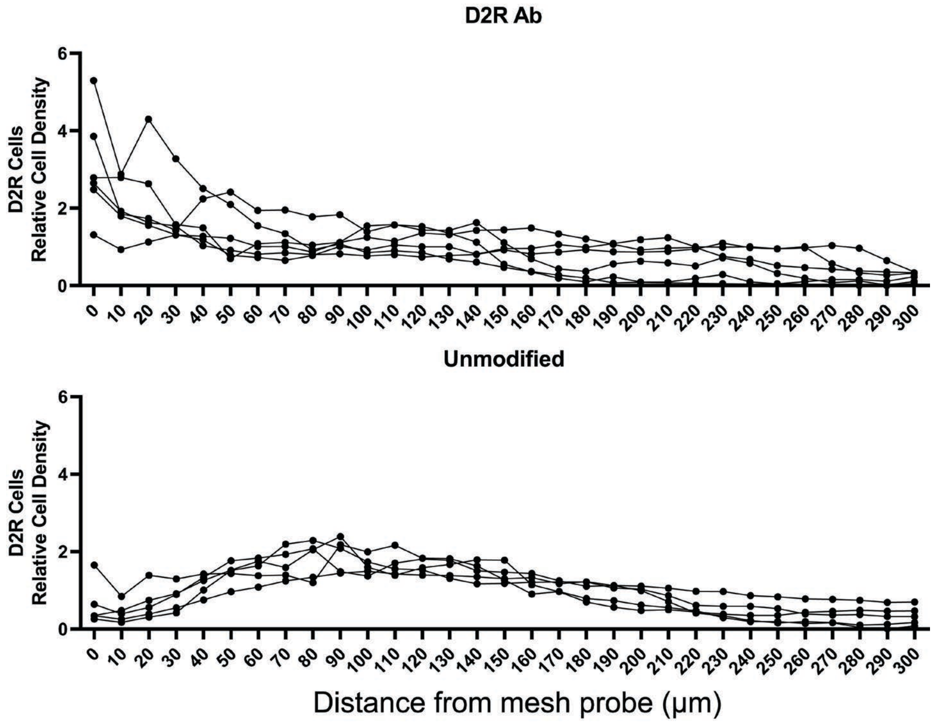


**Fig. S22.**

**Cell density distribution plots for individual samples in Fig. 4C.** Normalized cell density for D2R neurons in the CA1 region as a function of distance from surface of D2R Ab-modified (top) and unmodified (bottom) mesh probes. All D2R Ab-modified probes showed an increased association of D2R neurons, compared to the unmodified probes.

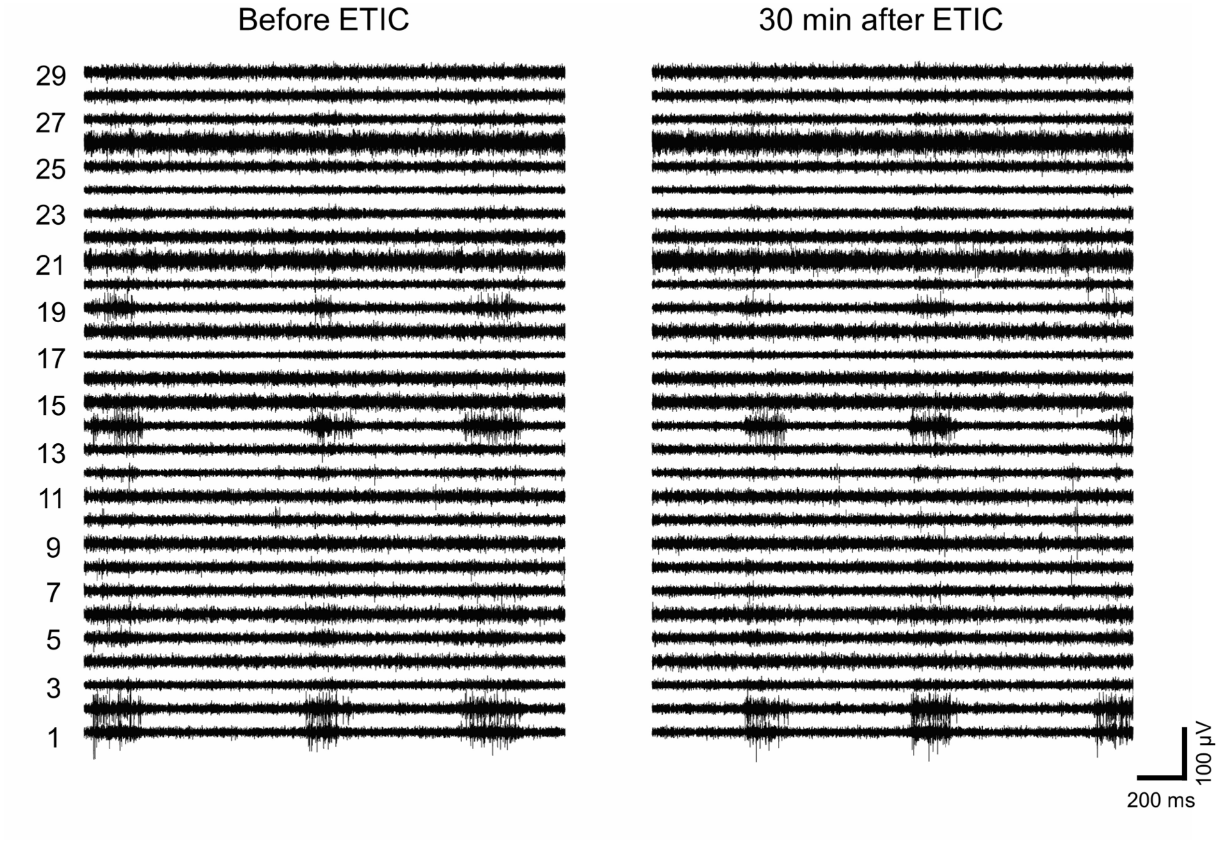


**Fig. S23.**

**Representative single-unit spike traces under anesthesia from an unmodified probe immediately before and 30 min after ETIC injection.** The x and y axes represent recording time and voltage. Channels with smaller numbers are implanted deeper. No clear difference in recording was observed before and after ETIC injection.

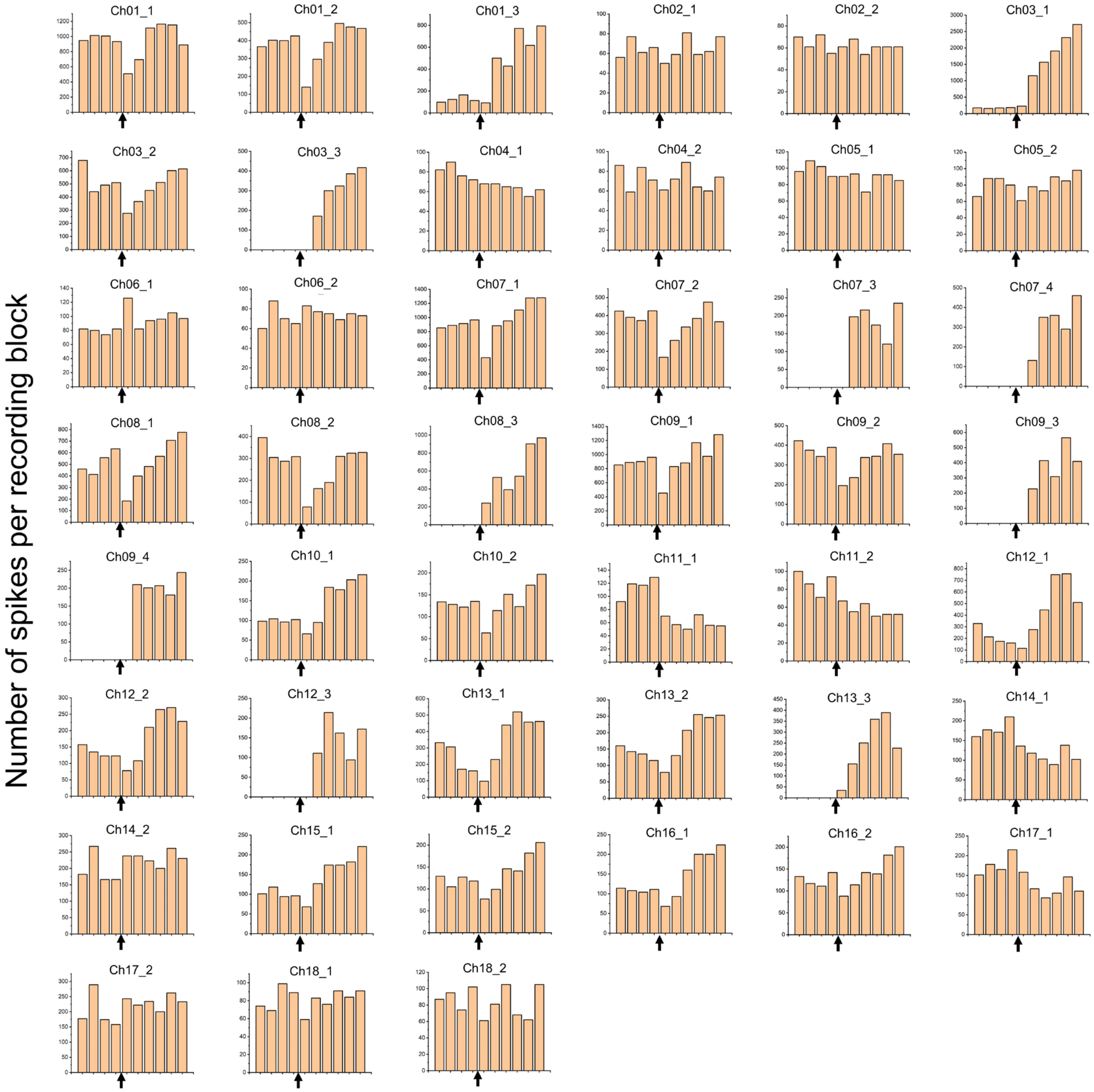


**Fig. S24.**

**Representative firing pattern changes of neurons recorded by the D2R Ab modified probe shown in Fig. 4D.** Neural activity was recorded from mice under stable anesthesia for 20 min before and 30 min after ETIC injection. Each bar represents the number of spikes recorded within a 5-min recording block. The black arrows denote the time point of ETIC injection. Many of the recorded neurons showed increased firing rate after ETIC injection.

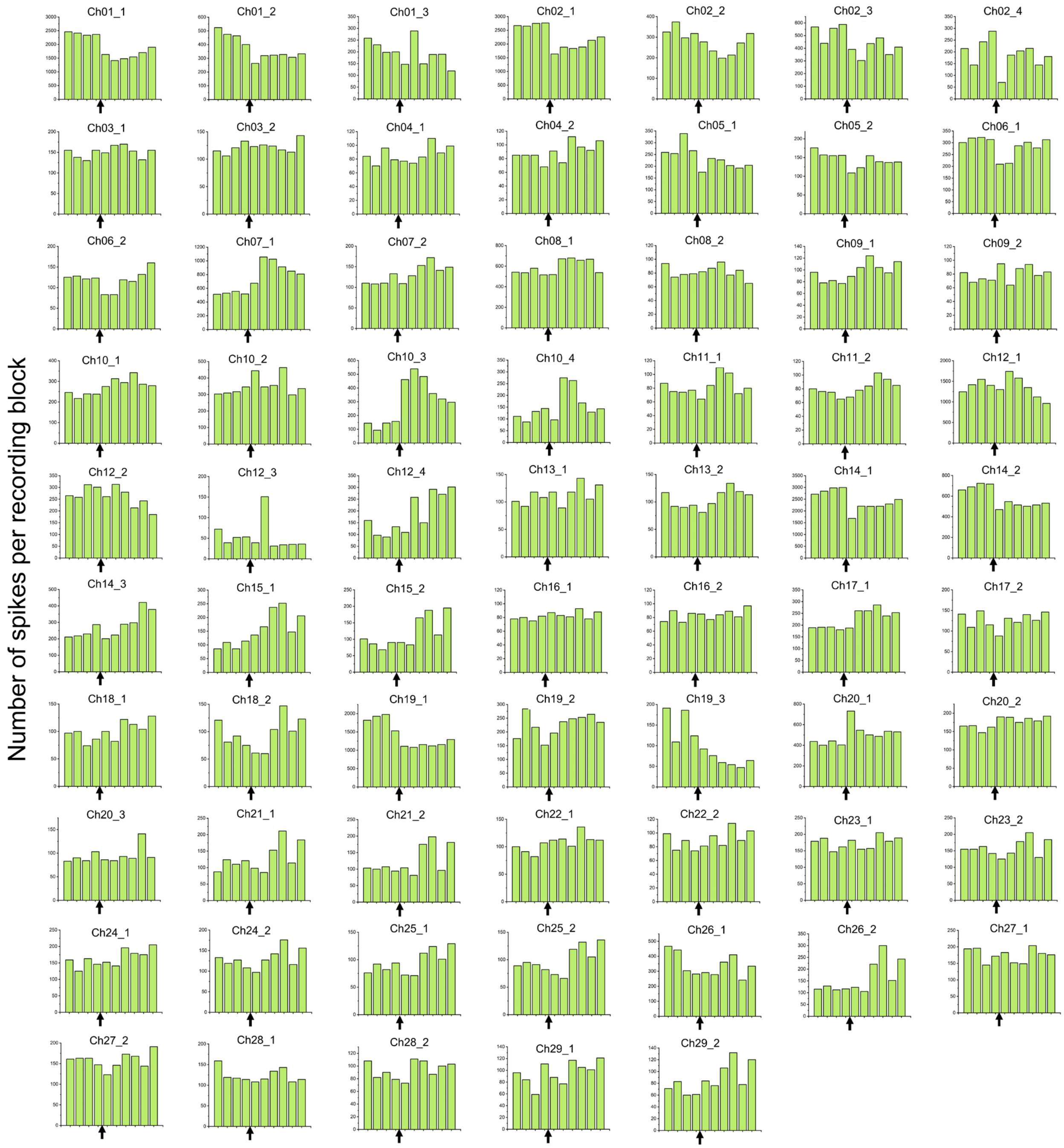


**Fig. S25.**

**Representative firing pattern changes of neurons recorded by the unmodified probe shown in fig. S23.** Neural activity was recorded from mice under stable anesthesia for 20 min before and 30 min after ETIC injection. Each bar represents the number of spikes recorded within a 5-min recording block. The black arrows denote the time point of ETIC injection. The majority of recorded neurons did not show changes in firing rate in response to ETIC injection.

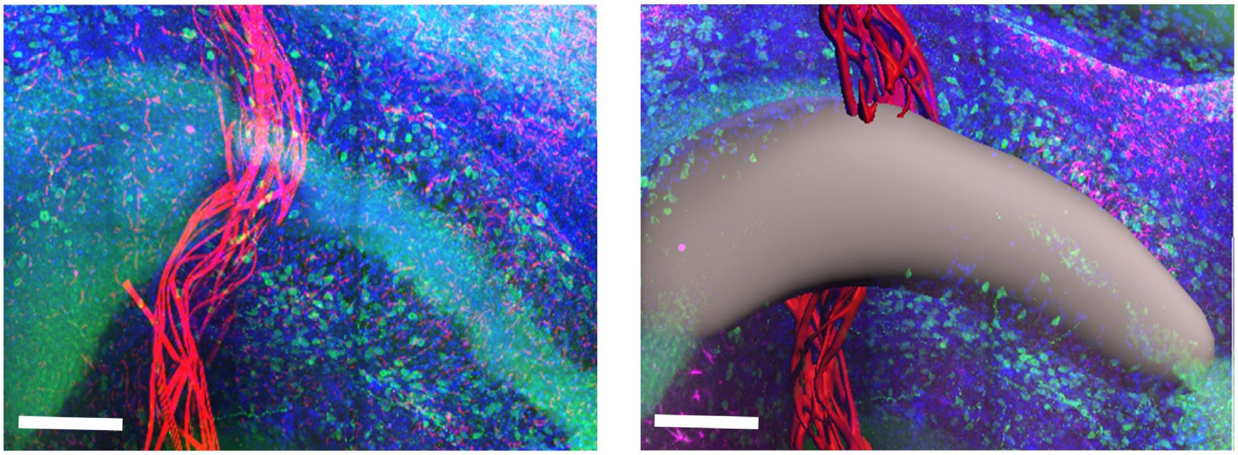

**Fig. S26.**

**Example of volumes drawn for quantification of the fluorescence intensity.** Regions were designated by hand using the boundaries of the CA1 (grey) or automatically via Imaris software edge detection for the mesh probe (red). Scale bars, 200 µm.
